# Structural mechanism of *LIN28B* nucleosome targeting by OCT4 for pluripotency

**DOI:** 10.1101/2023.01.03.522631

**Authors:** Ruifang Guan, Tengfei Lian, Bing-Rui Zhou, Yawen Bai

## Abstract

Pioneer transcription factors are essential for cell fate changes by targeting closed chromatin. OCT4 is a crucial pioneer factor that can induce cell reprogramming. However, the structural basis of how pioneer factors recognize the *in vivo* nucleosomal DNA targets is unknown. Here, we determine the high-resolution structures of the nucleosome containing human *LIN28B* DNA and its complexes with the OCT4 DNA binding region. Three OCT4s bind the pre-positioned nucleosome by recognizing non-canonical DNA motifs. Two use their POUS domains by forming extensive hydrogen bonds. The other uses the POUS-loop-POUHD region; POUHD serves as a wedge to unwrap ∼25 base pair DNA. Biochemical studies suggest that multiple OCT4s cooperatively open the H1-condensed nucleosome array containing the *LIN28B* nucleosome. Our study suggests a mechanism whereby OCT4s target the *LIN28B* nucleosome by forming multivalent interactions with nucleosomal motifs, unwrapping nucleosomal DNA, evicting H1, and cooperatively open closed chromatin to initiate cell reprogramming.

## INTRODUCTION

Gene regulation plays a crucial role in cell fate control. However, in eukaryotic cells, genomic DNA is packaged into chromatin by associating core and linker histones to form nucleosomes and chromatosomes (Luger et al., 1997; Zhou et al., 2021). Together with the compaction of chromatin to higher-order structures, a significant fraction of the DNA surface is not accessible for transcription factors. Nevertheless, a special group of transcription factors, termed pioneer factors, can recognize the closed chromatin sites (DNase I resistant) at enhancers (Iwafuchi-Doi and Zaret, 2014; Zaret, 2020). They facilitate the subsequent recruitment of other transcription factors to regulate gene expression and play critical roles in cell differentiation and development (Larson et al., 2021; Balsalobre and Drouin, 2022). For example, the forced expression of transcription factors OCT4, SOX2, KLF4, and c-MYC can induce pluripotent stem cells from somatic cells by accessing closed chromatin sites (Takahashi and Yamanaka, 2006; Soufi et al., 2012; Soufi et al., 2015). Functional genomics and biochemical studies reveal that OCT4 binds the nucleosome containing the human enhancer *LIN28B* DNA (162 bp, Figure 1A) during the reprogramming of fibroblasts to pluripotent cells (Kelly et al., 2012; Shyh-Chang et al., 2013; Soufi *et al*., 2015; Yu et al., 2007). The *LIN28B* locus is important for pluripotency reprogramming and OCT4 binding to the *LIN28B* nucleosome precedes *LIN28B* gene activation, which is silent in human fibroblasts and remains silent after 48-hour induction of OCT4, SOX2, KLF4, and c-MYC (Soufi *et al*., 2012; Soufi *et al*., 2015).

**Figure 1.**
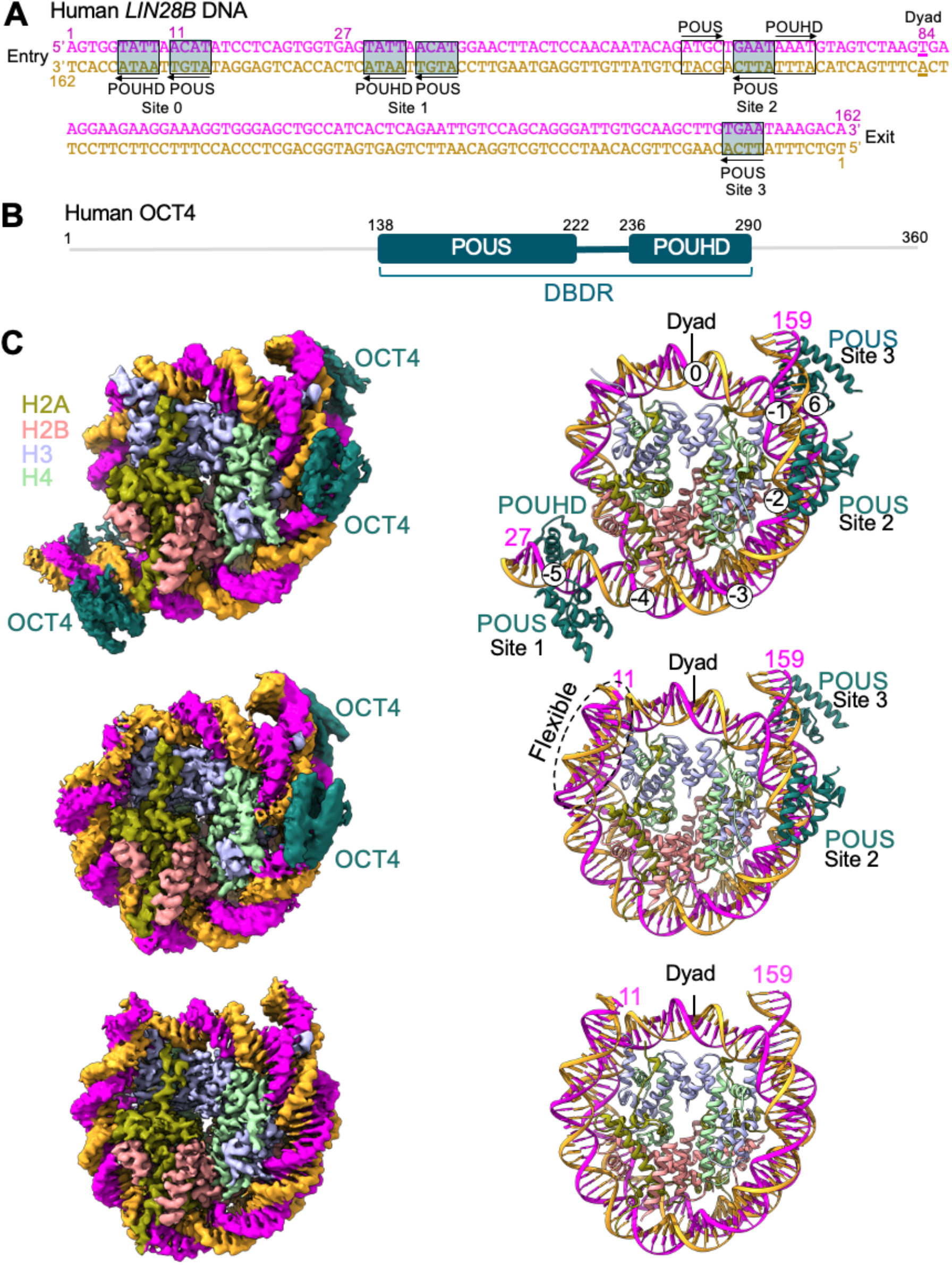
Overall structures of the *LIN28B* nucleosome bound to ^MBP-DBDR^OCT4. (A) The *LIN28B* DNA sequence. The boxes with transparent shade in cyan show the noncanonical DNA motifs for POUS and POUHD. The open boxes show the canonical motifs for POUS and POUHD, respectively. Arrows indicate the direction of the motifs. The base pair at the dyad is highlighted with the underline. (B) Diagram illustration of the domain organization of human OCT4 protein with the residue numbers that mark the boundaries of the domains. (C) Density maps (left) of the *LIN28B* nucleosome bound to three (top), two (middle) and 0 ^MBP-DBDR^OCT4s and their corresponding structural models (right). The two DNA strands are in magenta and orange, respectively. See also Figures S1-3 and Table S1 and Video S1.

To understand how OCT4 recognizes the *LIN28B* nucleosome, we want to solve the structure of the nucleosome bound to OCT4. However, determining the high-resolution structures of the nucleosomes containing genomic DNA sequences and their complexes with the pioneer factors is technically challenging. Genomic nucleosomes are fragile and pioneer factors tend to dissociate during sample preparation for structural studies by single particle cryo-electron microscopy (cryo-EM). So far, our knowledge of the interactions between pioneer factors and nucleosomal DNA comes from studies using model systems with non-genomic DNA sequences that bind the core histones tightly. These earlier studies have led to various binding modes of pioneer transcription factors (Zhu et al., 2018; Donovan et al., 2019; Michael et al., 2020; Dodonova et al., 2020; Tanaka et al., 2020; Guan et al., 2021). In the case of OCT4, it only uses the POUS domain to recognize its canonical DNA motif (ATGC) that is incorporated into the Widom 601 (W601) DNA near the entry-exit site of the nucleosome (Michael *et al*., 2020). Other biochemical and modeling studies suggest that OCT4 may also use the POUS domain to recognize the canonical motif in the inner region of the nucleosomal DNA (Soufi *et al*., 2015; Roberts et al., 2021). Together, these studies lead to the current view that OCT4 uses the POUS domain to recognize its partial canonical motif in the nucleosomal DNA. However, functional studies have shown that the entire DBD region (DBDR) of OCT4 (POUS-loop-POUHD; residues 138-290, Figure 1B), termed ^DBDR^OCT4, is essential for generating induced pluripotent stem cells (Jin et al., 2016; Kong et al., 2015; Roberts *et al*., 2021).

Here, we report the cryo-EM structures of the nucleosome containing human *LIN28B* DNA derived from the chromatin locus bound by OCT4 during cell reprogramming and its complexes with the OCT4 POUS-loop-POUHD region. Together with biochemical experiments, these structures suggest that multiple OCT4 molecules can target non-canonical DNA motifs in the *LIN28B* nucleosome, which leads to the eviction of linker histone H1 and unwrapping of the nucleosomal DNA, providing insights into how OCT4 opens closed chromatin.

## RESULTS

### Structural determination of the *LIN28B* nucleosome bound to ^MBP-DBDR^OCT4

Previous biochemical studies have shown that the full-length OCT4 binds the *LIN28B* nucleosome with ∼1:1 stoichiometry in the electrophoretic mobility shift assay (EMSA) (Soufi *et al*., 2015; Echigoya et al., 2020; Roberts *et al*., 2021). We tried to reconstitute the *LIN28B* nucleosome-OCT4 complex but found that the full-length OCT4 showed very low solubility and tended to aggregate when bound to the nucleosome. To improve the solubility of OCT4, we fused maltose-binding protein (MBP) to the N-terminus of the full-length OCT4 and ^DBDR^OCT4 through a flexible linker, referred to ^MBP^OCT4 and ^MBP-DBDR^OCT4, respectively. MBP has a chaperone function that helps solubilize recombinant proteins but does not bind the nucleosome (Chittori et al., 2018; Kapust and Waugh, 1999). Indeed, MBP-fused OCT4 proteins are substantially more soluble. Unexpectedly, EMSA experiments showed that multiple ^MBP-DBDR^OCT4, ^DBDR^OCT4, and ^MBP^OCT4 molecules could bind the nucleosome with the same apparent affinity, with ^MBP-DBDR^OCT4 displaying higher solubility (Figure S1). Accordingly, using the single-chain antibody (scFv)-aided cryo-electron microscopy (cryo-EM) approach that helps prevent nucleosome dissociation (Zhou et al., 2019), we collected images of the nucleosome bound to scFv without and with ^MBP-DBDR^OCT4. We obtained density maps of nucleosome-scFv_2_, nucleosome-scFv_2_-(^MBP-DBDR^OCT4)_3_, and nucleosome-scFv_2_-(^MBP-DBDR^OCT4)_2_ at overall resolutions of ∼2.6 Å and built the structural models accordingly (Figure 1C, Figures S2 and S3, Tables S1 and S2, Video S1).

### Overall structures of the *LIN28B* nucleosome and its complexes with OCT4

In these structures, all nucleosomes are uniquely positioned with the same dyad location (Figure 1A). scFv binds on the core histone surface in the nucleosome, showing no interactions with DNA and ^MBP-DBDR^OCT4 (Figures S2 and S3). The structure of the free 162 bp *LIN28B* nucleosome without ^MBP-DBDR^OCT4 includes well-defined 149 bp DNA. In the nucleosome-scFv_2_-(^MBP-DBDR^OCT4)_3_ complex, ^MBP-DBDR^OCT4s bind the noncanonical DNA motifs that are in the reverse direction (relative to the canonical motifs) (Figure 1A) at three sites near the super-helical locations (SHL) -5, -1.5, and +6.5, respectively. Hereafter, we refer to the motifs in nucleosomal DNA as nucleosomal motifs (Nu-motifs) to distinguish them from those in free DNA.

At site 1, the whole ^DBDR^OCT4 binds the Nu-motif; POUHD is wedged between the DNA and the nucleosome core, unwrapping ∼25 bp nucleosomal DNA from the entry side of the nucleosome. At sites 2 and 3, only POUS recognizes the Nu-motifs; the loop-POUHD region is missing, presumably flexible. In the nucleosome-scFv_2_-(^MBP-DBDR^OCT4)_2_ complex, POUS binds at sites 2 and 3 like those in the nucleosome-scFv_2_-(^MBP-DBDR^OCT4)_3_ complex. We observed weaker and broad density for ∼15 bp DNA at the entry site of the nucleosome (Figure S3J), indicative of flexible conformation, likely caused by ^MBP-DBDR^OCT4 binding.

### Specific OCT4 binding at sites 1-3

It is unexpected that OCT4 binds the non-canonical motifs at all three sites rather than the canonical motifs. To confirm that OCT4 binding at sites 1-3 is sequence-specific, we used the non-canonical DNA motifs at sites 1-3 in the *LIN28B* nucleosome to substitute the nucleotides at the corresponding locations in the W601 nucleosome to obtain a mutant nucleosome, ^W601_3sites^nucleosome. EMSA titration experiments of ^MBP-DBDR^OCT4 binding showed four sharp bands above the free ^W601_3sites^nucleosome (Figure 2A), suggesting that ^MBP-DBDR^OCT4 can bind the nucleosomal motifs specifically. The control experiment showed that ^MBP-DBDR^OCT4 binding to the free W601 nucleosome leads to one sharp band above the free nucleosome (Figure 2B), suggesting a specific binding site for OCT4 exists in the W601 nucleosomal DNA. Therefore, ^W601_3sites^nucleosome consists of three specific binding sites from sites 1-3 and one site from the W601 nucleosome. Quantitative analysis showed that ^MBP-DBDR^OCT4 binds ^W601_3sites^nucleosome much stronger than the W601 nucleosome but about two times weaker than the 162 bp *LIN28B* nucleosome (Figure 2C). These results provide additional evidence to support that OCT4 binding to the *LIN28B* nucleosome at sites 1-3 is sequence-specific and suggests that there is a fourth binding site in the 162 bp *LIN28B* nucleosome (see later results).

**Figure 2.**
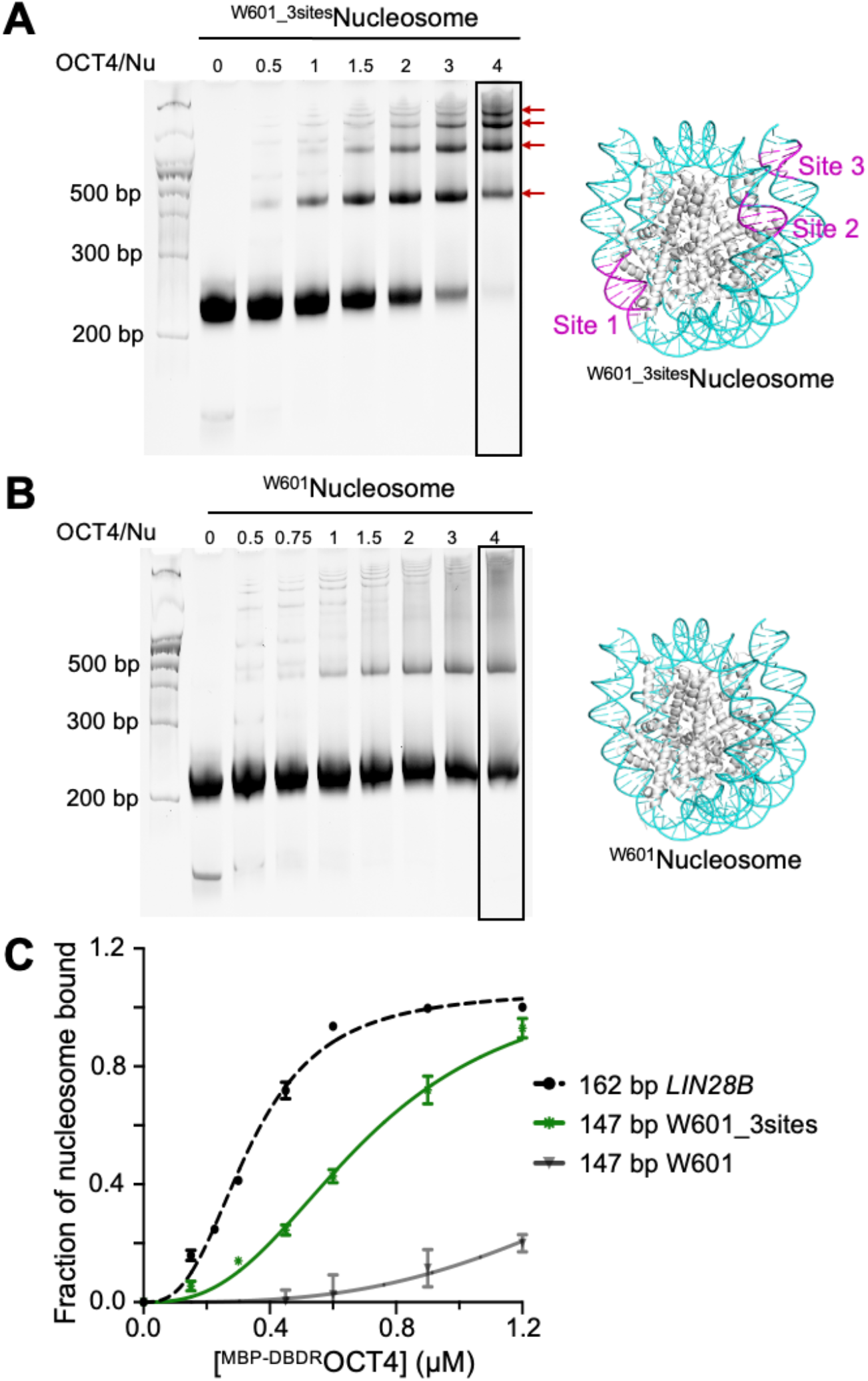
OCT4 binding at sites 1-3 are sequence specific. (A) EMSA showing the binding of ^MBP-DBDR^OCT4 (OCT4) to the ^W601-3sites^nucleosome (Nu) (left). The red arrows indicate the sharp bands. The locations of three sites are shown in magenta in the W601 nucleosome structure (right). (B) EMSA showing the binding of ^MBP-DBDR^OCT4 (OCT4) to the ^W601^nucleosome (Nu) (left) with the W601 nucleosome structure (right). (C) Plots showing the bindings of ^MBP-DBDR^OCT4 to the nucleosomes consisting of *LIN28B* W601-3sites, and W601 DNA. The error bars represent standard deviations from three independent experiments. Apparent Kd values were obtained by fitting the data to Hill equation: 0.34 ± 0.04 µM and 0.71 ± 0.04 µM for the *LIN28B* nucleosome and the ^W601-3sites^nucleosome, respectively. The data for the W601 nucleosome could not be fit reliably due to very weak binding. See also Figure S1.

### Interactions between ^DBDR^OCT4 and the Nu-motifs

At site 1, ^DBDR^OCT4 binds the Nu-motifs like the corresponding region of human OCT1 binding to similar DNA motifs in a DNA fragment (Figure 3A-C and Figure S4). Notably, the fourth base pairs, different in the corresponding motifs, do not form hydrogen bonds with the DBDs. The loop region between POUS and POUHD, consisting of three Arg and two Lys residues (RK motif, residues 230-234), forms a basic patch that interacts with the DNA with a narrow minor groove consisting of an AT-track (Figure 3A and Figure S5) (Kong *et al*., 2015). We found that mutation of the basic residues to Ala reduces the binding affinity and specificity between ^MBP-DBDR^OCT4 and the nucleosome, consistent with the earlier study that removing the positively charged residues not only disrupts their interactions with the DNA but also alters the structure of ^DBDR^OCT4 (Figure S5B, C) (Kong *et al*., 2015).

**Figure 3.**
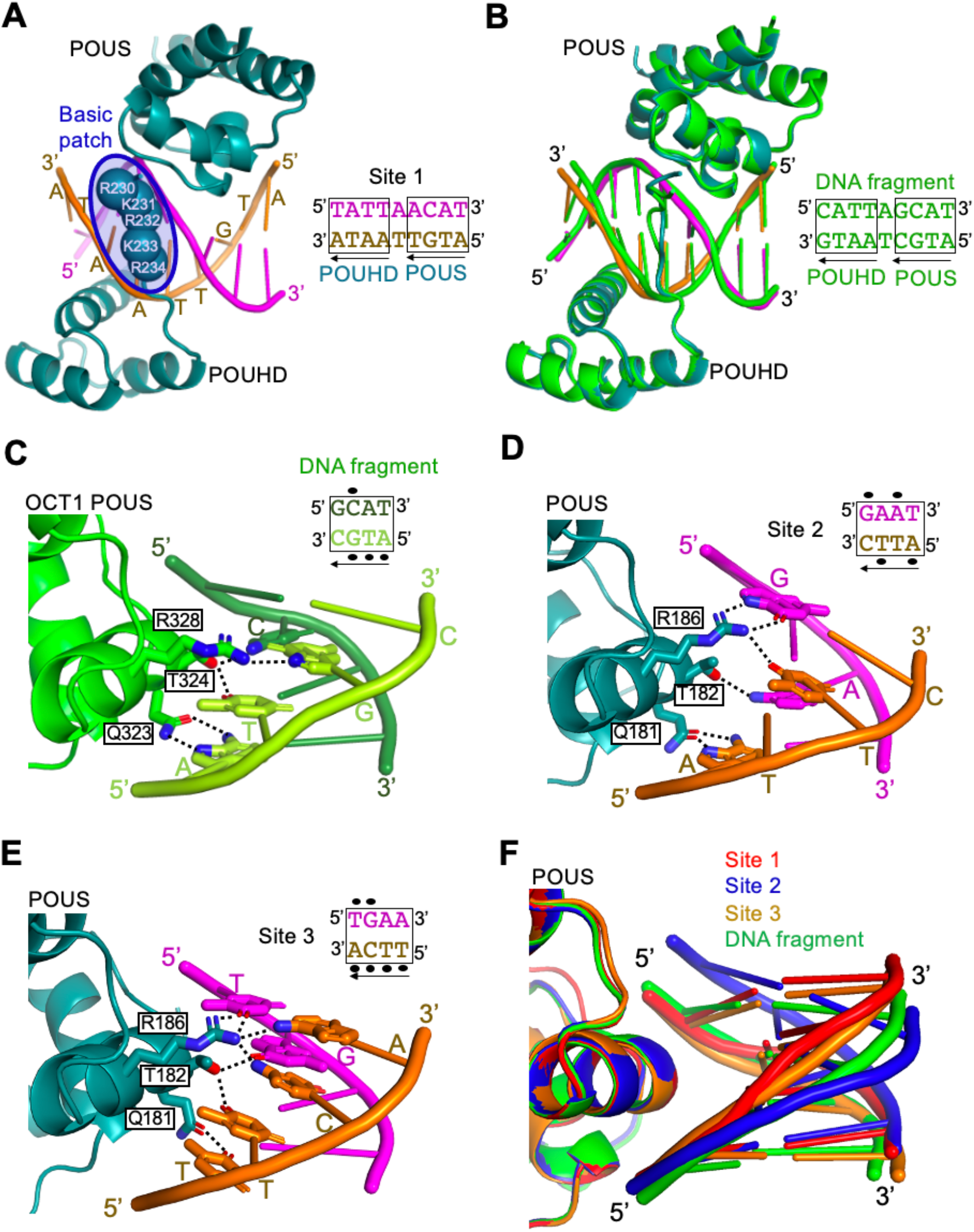
Interactions between OCT4 and the noncanonical motifs. (A) Structure of ^MBP-DBDR^OCT4 bound to Nu-motif at site 1. The basic patch residues that interact with the minor groove of DNA are represented with spheres centered at the backbone Cα carbon atoms. (B) Alignment of the structures of ^MBP-DBDR^OCT4 bound to the Nu-motif at site 1 in the nucleosome and human OCT1 DBDR bound to the DNA fragment in the crystal structure (PDB ID: 1GT0). (C) Illustration of hydrogen bonds (dashed lines) formed between POUS and the canonical DNA motif (ATGC) in the DNA fragment (PDB ID: 1GT0). They were drawn based on the distance (< 3.2 Å) between the two polar atoms. The side chains are shown in sticks. The nitrogen and oxygen atoms involved in the hydrogen bond formation are colored in blue and red, respectively. The black ovals close to the bases of the motifs indicate their involvement in hydrogen bond formation. (D) Illustration of the hydrogen bonds (dashed lines) between POUS and the Nu-motif at sites 2. (E) Illustration of the hydrogen bonds (dashed lines) between POUS and the Nu-motifs at site 3. They may not form simultaneously due to angle restriction. (F) Comparison of the conformations of POUS bound Nu-motifs at sites 1-3 and the canonical motif in the DNA fragment. The structures are aligned on the POUS domains. See also Figures S4 and S5.

At sites 2 and 3, POUS Arg186 forms multiple hydrogen bonds with the bases in the Nu-motifs’ third and fourth base pairs (Figure 3D, E, and Figure S4). In contrast, only one hydrogen bond forms between the corresponding Arg residue in POUS and the third base of the canonical motif (ATGC) in the DNA fragment (Figure 3C). Notably, the DNA shape in the nucleosome at site 2 differs substantially from those in sites 1, 3, and the DNA fragment (Figure 3F). In addition, DNA bending by the core histones at site 2 enlarges the major groove. These results reveal that POUS Arg186, with a long side chain and the guanidino group, can form multiple hydrogen bonds with the Nu-motifs with non-canonical DNA sequence. The lack of AT-tracks at sites 2 and 3 is likely the cause for the missing of the basic patch (Figure S5A). The formation of multiple hydrogen bonds between POUS and the Nu-motifs provides additional evidence that the binding of the non-canonical motifs by ^DBDR^OCT4 at sites 1-3 are sequence-specific.

### Distinct binding modes of OCT4

In the nucleosome-scFv_2_-(^MBP-DBDR^OCT4)_3_ structure, ^MBP-DBDR^OCT4 binding leads to unwrapping of nucleosomal DNA and the relocation of histone H2B N-terminal tail between the two DNA gyres near site 1 (Figure 4A). The DNA conformations at sites 2 and 3 remain unchanged compared to those in the nucleosome without ^MBP-DBDR^OCT4 (Figure 4B). To understand why the canonical motifs, close to site 2 (Figure 1A), in the *LIN28B* nucleosome are not recognized by OCT4, we aligned the canonical motif (ATGC or AAAT) in the nucleosome structure to the corresponding motif in the DNA fragment bound to POUS or POUHD. Both POUS and POUHD clash with the core histones (Figure 4C), suggesting that the intrinsic positioning of the *LIN28B* nucleosome plays a determinant role in decoupling OCT4 recognition of the nucleosomal motifs from those in the free DNA (Roberts *et al*., 2021). Another unexpected finding is that the modes of OCT4 binding to the *LIN28B* nucleosome are distinct in comparison with those identified in the previous study using the W601 nucleosome as the host with the incorporation of the canonical motif as the guest (Figure 4D, E) (Michael *et al*., 2020). In our structures, POUS binds the major grooves at super-helical locations (SHL) -4.5, -1.5, and 6.5, and POUHD serves as a wedge to unwrap nucleosomal DNA. In contrast, in the W601-nucleosome-based structure, POUS binds at either SHL -5.5 or +5.5, and POUHD is missing.

**Figure 4.**
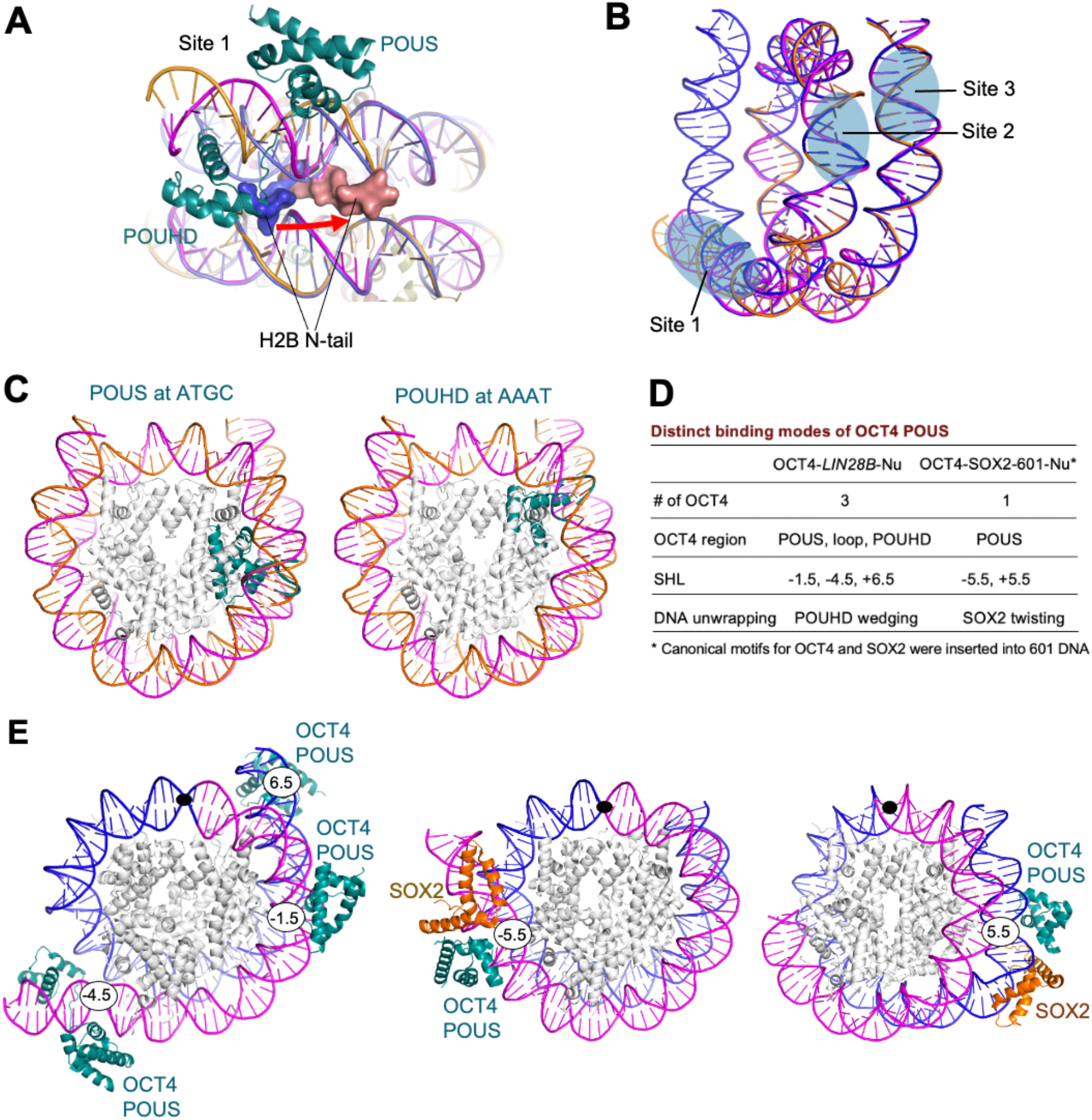
Conformational changes and distinct binding modes of OCT4. (A) The N-terminal tail of H2B in the free nucleosome (blue) near site 1 relocates to a new position (orange) when site 1 is bound by OCT4. (B) Overlay of the nucleosomal DNA in the nucleosome-scFv_2_-(^MBP-DBDR^OCT4)_3_ complex (orange) and the nucleosome (blue) with the highlighted sites 1-3. (C) Modeling of OCT4 POUS and POUHD binding to their canonical motifs (ATGC and AAAT) in the *LIN28B* nucleosome. Each motif in the nucleosome is aligned with the corresponding motif in the crystal structure of the DNA fragment bound to either POUS or POUHD. (D) Summary of the differences in OCT4 binding modes between our structure and the two structures in the W601-based nucleosome (see E). (E) Comparison of our structure (left) and the two structures of W601-based nucleosomes (PDB IDs: 6T90 and 6YOV) (middle and right). The filled ovals indicate the nucleosome dyad. The numbers are the SHLs. See also Figure S3.

### Unwrapping nucleosomal DNA by OCT4

Our structures suggest that OCT4 binding can lead to the unwrapping of the nucleosomal DNA at the entry side of the *LIN28B* nucleosome (Figure 5A). The conformational change between the free nucleosome and that bound to OCT4 would lead to differences in sensitivity to micrococcal nuclease (MNase) and the distance between the entry side DNA and core histones. To verify it, we performed micrococcal nuclease (MNase) digestion and fluorescence resonance energy transfer (FRET) experiments by titrating the nucleosome with ^MBP-DBDR^OCT4. As the ratio of ^MBP-DBDR^OCT4 to nucleosome increased, we found that MNase cut the DNA in the *LIN28B* nucleosome more efficiently (Figure 5B). The FRET signal, with donor (Cy3) at the entry site of the DNA and the acceptor (Cy5) at histone H2A residue 116, respectively, became weaker (Figure 5C). In contrast, when the donor is at the exit site, OCT4 binding does not cause change of the FRET signal (Figure S6). These results indicate that OCT4 only unwarps the nucleosomal DNA at the entry site, in agreement with our structures (Figure 1C).

**Figure 5.**
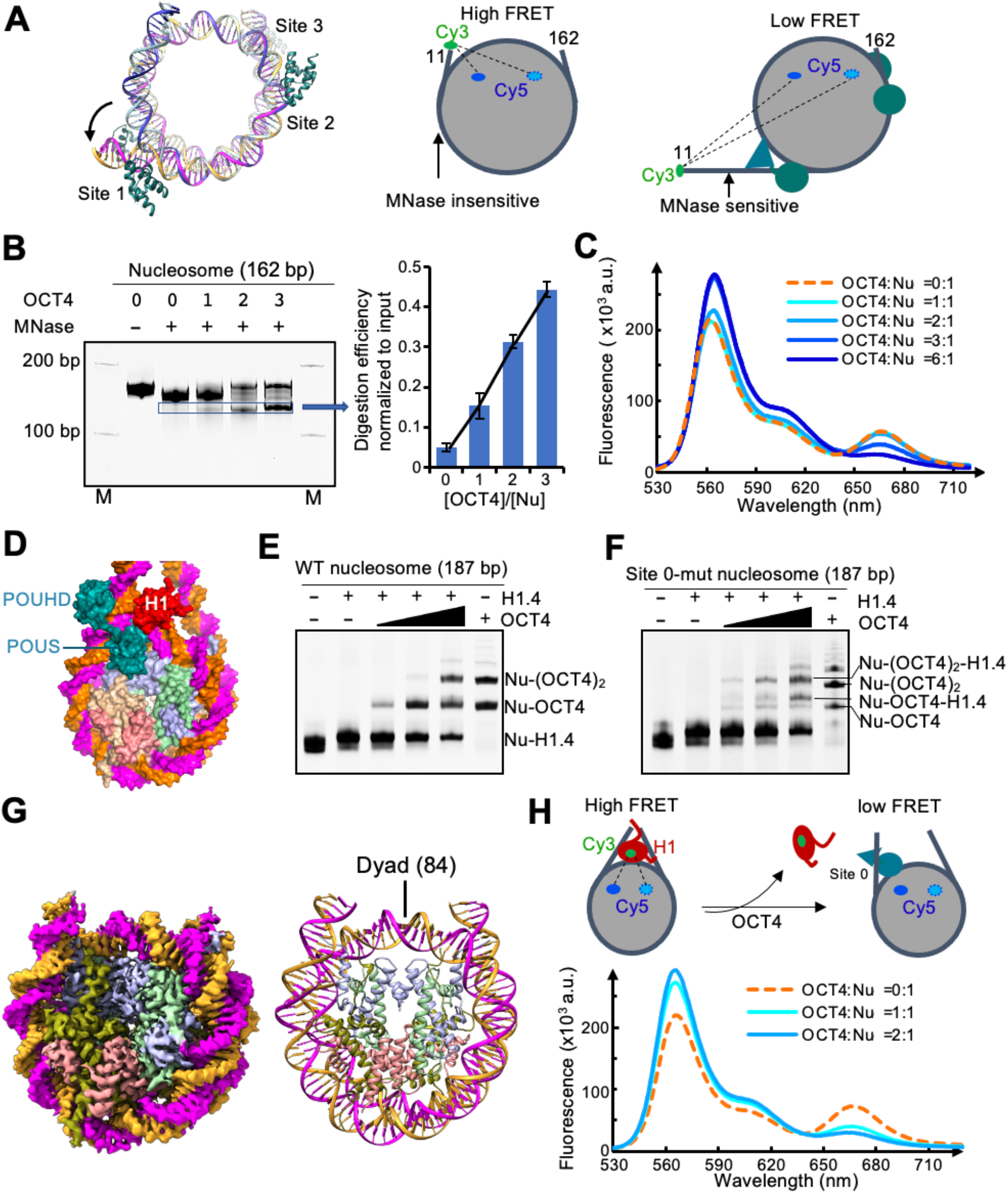
Biochemical studies of DNA unwrapping, site 0 binding, and H1 eviction. (A) Comparison of the DNA structures in the *LIN28B* nucleosome alone (blue) and that bound to three ^MBP-DBDR^OCT4s (magenta and orange) (left) and their anticipated sensitivity to MNase and FRET signals (middle and right). (B) MNase digestion of the *LIN28B* nucleosome with increasing ^MBP-DBDR^OCT4 concentration (Left). The numbers indicate the ratio of ^MBP-DBDR^OCT4 over the nucleosome. Quantification of the DNA fragment over the input (Right). Error bars represent standard deviations from three independent experiments. (C) FRET experiment with Cy3 labeled at the entry site of the DNA and Cy5 labeled at histone H2A residue 116. (D) Modeling of binding of ^DBDR^OCT4 to site 0 in the chromatosome by aligning the *LIN28B* nucleosome with the chromatosome structure (PDB ID: 7K5Y) using the core histones and by aligning the DBDR bound to the site 1 DNA to site 0. (E) EMSA for ^MBP-DBDR^OCT4 binding to the chromatosome. The concentrations of the nucleosome and H1.4 are 0.3 μM and 0.48 μM, respectively. [^MBP-DBDR^OCT4]/[nucleosome] is 0.25, 0.5, 1.0, or 2.0. (F) EMSA showing that binding of ^MBP-DBDR^OCT4 to the chromatosome containing site 0 mutation. The concentrations of the nucleosome and H1.4 are 0.3 and 0.48 μM, respectively. [^MBP-DBDR^OCT4]/[nucleosome] is 0.25, 0.5, 1.0, or 2.0. (G) Structure of the *LIN28B* nucleosome with 187 bp DNA and site 0 mutation, showing the same dyad as the 162 bp *LIN28B* nucleosome. (H) FRET signals of ^MBP-DBDR^OCT4 binding to the chromatosome containing Cy3 labeled on H1.4 at residue 26 and Cy5 at histone H2A residue 116. See also Figure S6.

### OCT4 binding at site 0 and eviction of H1

In the *LIN28B* DNA sequence (Figure 1A), another site (site 0) near the entry of the nucleosome has the identical DNA sequence as site 1. Structural modeling showed that OCT4 POUHD could bind the exposed motif at site 0 in the chromatosome, with POUS clashing with the linker histone globular domain and the DNA near the dyad (Figure 5D) (Zhou *et al*., 2021). This result suggests that OCT4 might bind site 0 and inhibit H1 binding, consistent with the earlier observation (Figure 2C) (Echigoya *et al*., 2020). To further verify it, we reconstituted the 187 bp *LIN28B* nucleosome without and with a site 0 mutation. We found that ^MBP-DBDR^OCT4 could largely replace linker histone H1 in the chromatosome at a 1:1 ratio of ^MBP-DBDR^OCT4 versus chromatosome in the EMSA experiments (Figure 5E). It failed to do so when site 0 was mutated to abolish ^MBP-DBDR^OCT4 binding at this site (Figure 5F). We solved the structure of the 187 bp nucleosome with site 0 mutation (Figures 5G and S1H-K). The dyad of the nucleosome is located at the same position as in the 162 bp nucleosome, indicating the extension of the linker DNA region and site 0 mutation do not cause repositioning of the nucleosome. We confirmed this conclusion by conducting the FRET experiments with Cy3 and Cy5 labeled on H1.4 residue 26 and H2A residue 116, respectively (Figure 5H). We found that at a 1:1 ratio of OCT4 over the chromatosome, FRET intensity decreased substantially.

### OCT4s cooperatively open H1-condensed nucleosome array

Structural modeling shows that four partial motifs in the *LIN28B* chromatosome are accessible to OCT4 (Figure 6A). To test if binding of OCT4 to the *LIN28B* nucleosome can open the closed chromatin, we reconstituted a H1-condensed nucleosome array consisting of 12 nucleosomes with the nucleosome repeat length of 197 bp. In this array, the *LIN28B* nucleosome is surrounded by five and six nucleosomes consisting of W601 sequence. We found that ^DBDR^OCT4 enhanced the DNA digestion efficiency of the nucleosome array by a site-specific DNase (DdeI) that cut the DNA in the *LIN28B* nucleosome (Figure 6B and Figure S7). Intriguingly, the digestion efficiency showed little change when the molecular ratio of ^DBDR^OCT4 over the nucleosome array was less than two but increased sharply when the ratio is between two to four (Figure 6C). When the ratio was more than four, the additional OCT4 has little effect on digestion efficiency. The S-shaped curve of digestion efficiency versus the ratio of OCT4 over the nucleosome array suggests that four ^DBDR^OCT4 molecules work cooperatively to open the chromatin target. This result agrees well with the mono-nucleosome study that four OCT4 molecules can bind to the *LIN28B* nucleosome specifically.

**Figure 6.**
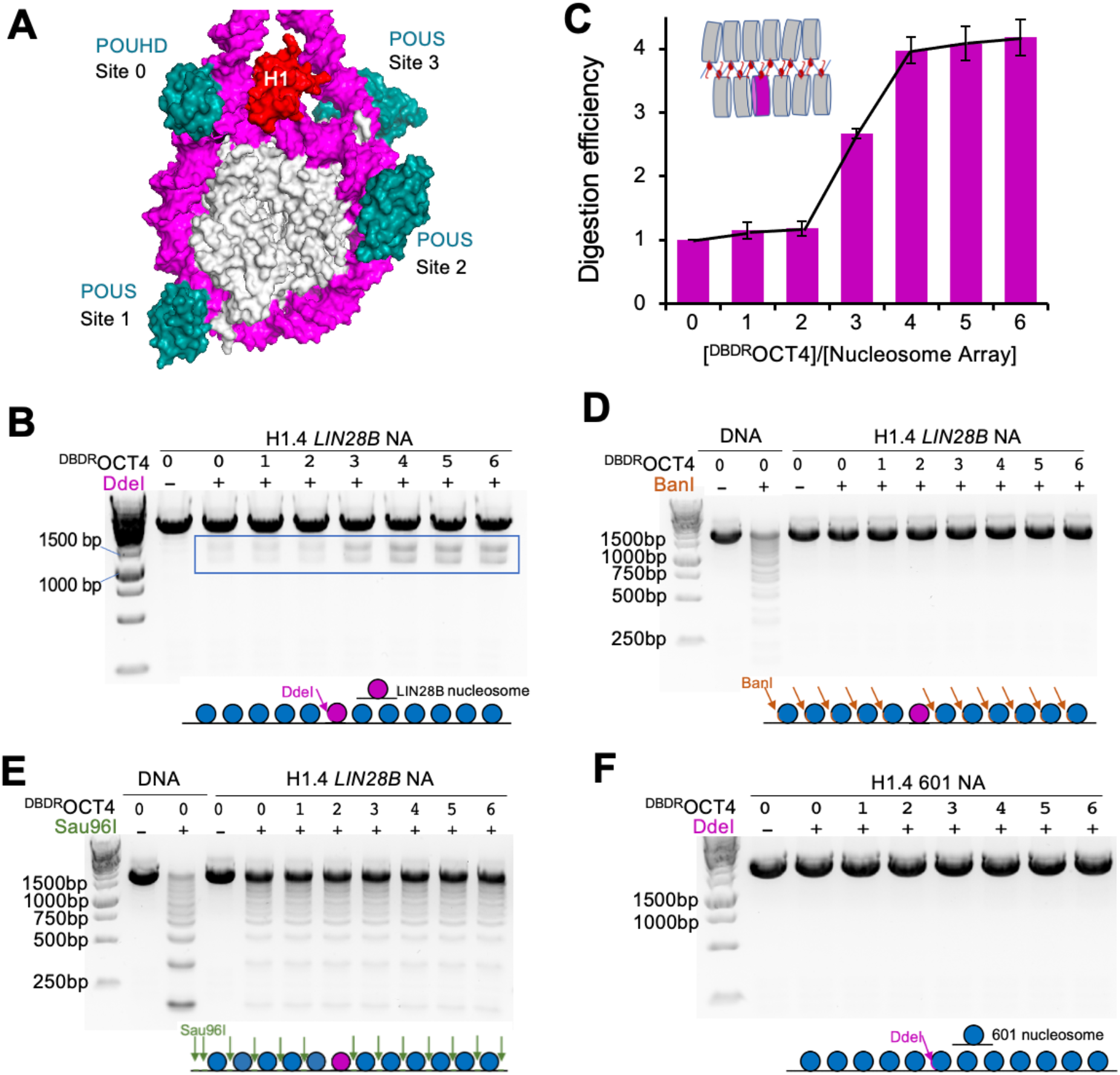
OCT4 opens H1-condensed nucleosome array containing the *LIN28B* nucleosome. (A) Illustration of four sites with partial motifs accessible to either POUS or POUHD in the *LIN28B* chromatosome. (B) Digestion of the nucleosome array by DdeI enzyme with increasing ratios of ^DBDR^OCT4 over the nucleosome array. (C) The relative digestion efficiency of the linker histone H1-condensed nucleosome array from (B). Error bars represent standard deviations from three independent experiments. (D) Digestion of the nucleosome array with BanI that cut the sites in the W601 nucleosome core. (E) Digestion of the nucleosome array with Sau96I that cut the sites in the linker DNA. (F) Digestion of the nucleosome array with the DdeI enzyme that cut the site within the W601 nucleosome that replaced the *LIN28B* nucleosome. See also Figure S7.

To further test whether OCT4 specifically binds to the *LIN28B* nucleosome in the nucleosome array and opens the nucleosome array locally or globally, we investigated the effect of OCT4 on the digestion efficiency of the nucleosome array with cutting sites at the DNA in all W601 nucleosome cores (Figure 6D), or all linker DNA (Figure 6E), or the W601 nucleosome that replaced the *LIN28B* nucleosome but keep the same cutting site (Figure 6F). We found OCT4 has no effects on the digestion efficiency in these cases. Therefore, these experiments indicate that OCT4 only opens the H1-condensed nucleosome array locally at the *LIN28B* nucleosome site.

## DISCUSSION

Our finding that multiple OCT4s recognize the *LIN28B* nucleosome through noncanonical DNA sequences using both POUS and POUHD is in contrast with the current view that only POUS binds at a single Nu-motif containing the canonical DNA sequence (Soufi *et al*., 2015; Michael *et al*., 2020; Roberts *et al*., 2021). We also find that OCT4 uses two linked domains to bind and block the DNA from rewrapping, explaining the mutation results that the entire DBDR is essential for initiating cell reprogramming (Jin *et al*., 2016; Kong *et al*., 2015; Roberts *et al*., 2021). By contrast, SOX11/SOX2 unwraps nucleosomal DNA through local DNA distortion (Dodonova *et al*., 2020). Thus, two DBDs, linked through a loop, working together to evict the linker histone and unwrap the nucleosome DNA is likely a unique feature of OCT4 for its pioneer function. It is worth noting that the previous study using the W601 nucleosome as the host shows that OCT4 loop-POUHD plays no role in either nucleosome recognition or nucleosomal DNA unwrapping (Michael *et al*., 2020). Computer simulation studies suggest nucleosome dynamics may play a role in determining the binding behavior of OCT4 as the DNA in the *LIN28B* nucleosome is significantly more dynamic than that in the W601 nucleosome (Huertas et al., 2020; Tan and Takada, 2020). Alternatively, SOX2 binding at the adjacent site might affect the binding of OCT4 (Figure 4E) (Michael *et al*., 2020). Together, these results suggest that OCT4 POUS can access broad locations in the nucleosomal DNA, and the nucleosome context plays a critical role in OCT4 binding.

Our study provides insights into the hypothesis of how pioneer transcription factors may target and open closed chromatin (Cirillo *et al*., 1998; Cirillo et al., 2002; Zaret, 2020). Structures and modeling show that four accessible Nu-motifs of POUS and POUHD exist in the *LIN28B* chromatosome (Figure 6A). However, chromatosomes in closed chromatin are likely to pack in a zigzagged manner (Figure S7G) (Garcia-Saez et al., 2018; Song et al., 2014), preventing direct access by OCT4. Nevertheless, transient exposure of the chromatosome could occur through intrinsic structural fluctuations (Zhou et al., 2018), allowing binding of multiple OCT4 molecules simultaneously, leading to cooperativity. For example, individual linker histones have a short residence time in condensed chromatin *in vivo* (Misteli et al., 2000). Notably, the sizes of the OCT4 DBDs and the globular domains of linker histones are comparable. As POUS can form multiple hydrogen bonds with the Nu-motifs, OCT4 may use POUS to scan the condensed chromatin for initial engagement. After the initial recognition of the transiently exposed nucleosome using partial Nu-motifs, OCT4 could further evict linker histones and partially unwrap the nucleosomal DNA (Figure 7). Our results suggest that this process requires OCT4s to work cooperatively.

**Figure 7.**
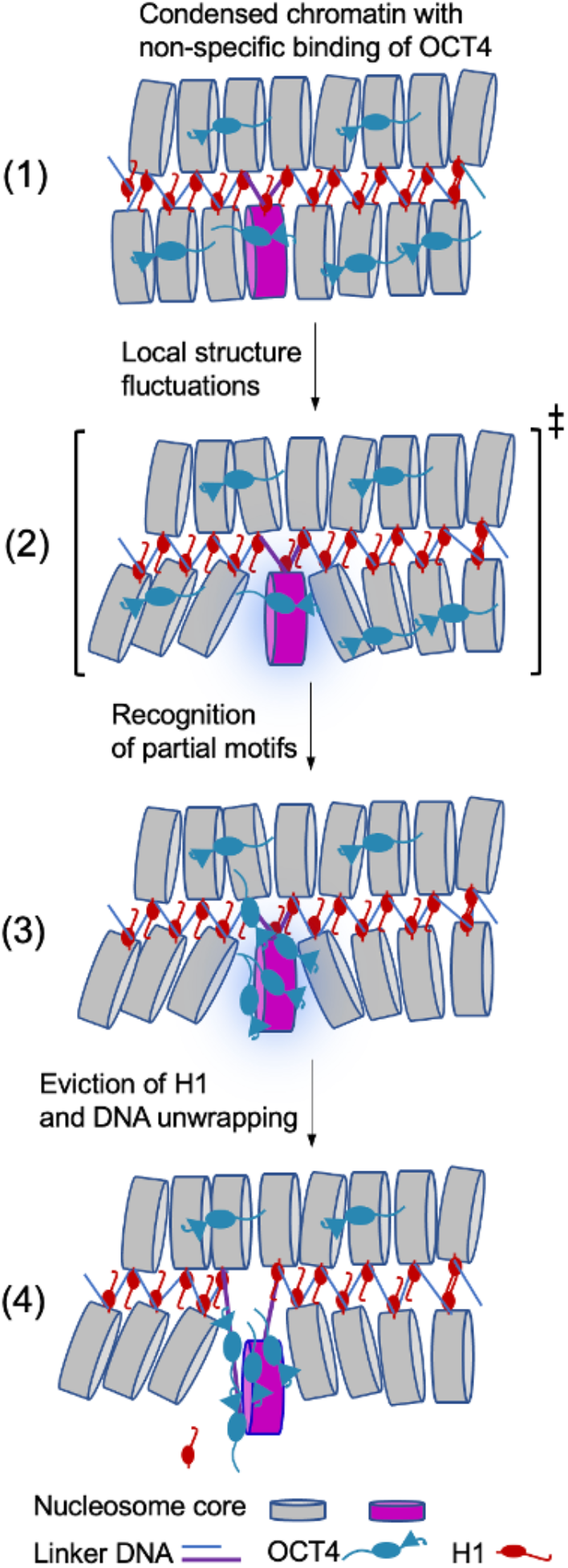
A working model for opening closed chromatin by OCT4. Diagram illustration of a model explaining the local opening of H1-condensed chromatin by OCT4. (1) Non-specific binding of OCT4 in searching for the *LIN28B* chromatosome. (2) Intrinsic stochastic fluctuations of the condensed chromatin structure transiently expose the *LIN28B* chromatosome. The parathesis with the ‡ indicate an activated high-energy state (analogues to the transition state in a chemical reaction or a high-energy intermediate state). (3) Recognition of the Nu-motifs by POUS and POUHD. (4) Eviction of linker histone H1, unwrapping of the nucleosomal DNA, and stabilization of the locally opened chromatin by OCT4.

The formation of the multi-valent binding of a pioneer transcription factor is likely to increase its overall time associated with the nucleosome through mass action (Tang et al., 2022), facilitating the recruitment of other transcription factors for transcription activation. For multi-valent binding to occur, however, it may require high concentration of the pioneer factor, which could be achieved through liquid-liquid phase separation (Alberti et al., 2019). Indeed, recent studies show OCT4 participates in liquid-liquid phase separation through its intrinsically disordered activation domain that promotes cell reprogramming (Boija et al., 2018; Wang et al., 2021).

Finally, we speculate that the general structural features for mammalian pioneer transcription factors to engage the chromatin may include multiple Nu-motifs, eviction of linker histones, and unwrapping of nucleosomal DNA. We anticipate our approach will inspire future investigations of other pioneer transcription factors bound to nucleosomes of their *in vivo* target (Cirillo et al., 1998; Fernandez Garcia et al., 2019), providing insights into broader biological functions.

### Limitations of the study

Our study provides structural insights into how OCT4 recognizes the *LIN28B* nucleosome and a working model for opening closed chromatin by OCT4 during cell reprogramming. We used the DNA binding region of OCT4 for structural study since it has a better solubility, which does not show whether the intrinsically disordered regions in the full-length protein interact with the nucleosome. Nevertheless, it is supported by the earlier structural study that the tails are not found to interact with the nucleosome (Michael *et al*., 2020), and ^DBDR^OCT4 and the full-length OCT4 show similar binding affinity (Figure S1). Also, this study does not provide the structural information of OCT4 bound to site 0 of the nucleosomal DNA and how OCT4 evicts the linker histone. In addition, how the four OCT4 molecules coordinate with each other to open the chromatin and how OCT4 facilitates the binding of SOX2, KLF4 and c-Myc to the *LIN28B* nucleosome remain to be investigated (Soufi *et al*., 2015). Our structure provides the platform for future study of these questions.

### Experimental Procedures

#### Expression and purification of histones

Recombinant human histones H2A, H2A^L116C^, H2B, H3, H4, H1.4, and H1.4 ^K26C^ were expressed individually in Escherichia coli BL21(DE3) cells as described in a previous study (Zhou *et al*., 2021). All mutations in our study were generated using QuikChange kit (Agilent). Briefly, E. coli cells harboring each histone expression plasmid were grown at 37 °C in 2 x YTB Broth. 0.3 mM IPTG was added to induce recombinant protein expression for 3 h at 37 °C, When OD_600_ reached around 0.6-0.8. The cells were harvested and resuspended in 50 ml of buffer A (50 mM Tris-HCl, 500 mM NaCl, 1 mM PMSF, 5% glycerol, pH 8.0), followed by sonication on ice for 60 min. The cell lysates were centrifuged at 35,000 RPM for 20 min at 4 ºC. The pellet containing histones was resuspended in 50 ml of buffer A and 7 M guanidine hydrochloride. The samples were rotated for 12 h, and the supernatant was recovered by centrifugation at 35,000 RPM for 60 min at 4 ºC. The supernatants were dialyzed against buffer C (5 mM Tris-HCl, pH 7.4, 2 mM 2-mercaptoethanol, 7 M urea) for three times. The supernatant was loaded to Hitrap S column chromatography (GE Healthcare). The column was washed with buffer D (20 mM sodium acetate, pH 5.2, 100 mM NaCl, 5 mM 2-mercaptoethanol, 1 mM EDTA, and 6 M urea). The histone protein was eluted by a linear gradient of 100 to 800 mM NaCl in buffer D. The purified histones were dialyzed against water for three times, and freeze-dried.

### Over-expression and purification of OCT4 proteins

The plasmid (pET30a) containing the human ^MBP-DBDR^OCT4 with a His-tag at the C-terminus and a Tobacco Etch Virus (TEV) recognition site (ENLYFQG/S) between MBP and OCT4 was cloned using NdeI and EcoRI restriction sites (Table S1). Ala mutants for the basic patch residues in the loop region including R232A/R234A, R232A/K233A/R234A, R230A/K231A/R232A/K233A/R234A, which are termed ^MBP-DBDR^OCT4^2A, MBP-DBDR^OCT4^3A, MBP-DBDR^OCT4^5A^, respectively. The plasmids harboring the above mutations were transformed into BL21(DE3) cells and grown in LB medium. When OD was about 0.8, 0.5 mM IPTG was added to induce protein expression at 18 ºC for 18 hrs. Cells were harvested and resuspended in the buffer containing 20 mM Tris pH 7.4, 1 M NaCl, 2 mM DTT. After sonification and ultracentrifuge, the supernatant went through Nickle affinity beads. The column was washed by the buffer containing 20 mM Tris pH 7.4, 1 M NaCl, 2 mM DTT, 50 mM imidazole, then eluted by the buffer containing 20 mM Tris pH 7.4, 1 M NaCl, 2 mM DTT, 300 mM imidazole. The solution containing the fusion protein was concentrated to ∼1 ml and went through gel filtration on Superdex 200 10/300 increase column (GE healthcare) equilibrated with buffer containing 20 mM Hepes pH 7.3, 500 mM NaCl, 2 mM DTT. The purified protein was concentrated to ∼40 µM and stored at -80 ºC. ^DBDR^OCT4 was obtained by cutting the ^MBP-DBDR^OCT4 with TEV protease (Genscript) for overnight at 4 ºC. The digested sample was mixed with Amylose resin (NEB) and incubated 3 hours. After centrifuge, the supernatant went through Nickle affinity beads. The beads were washed with the buffer containing 20 mM Hepes pH 7.4, 500 mM NaCl, 2 mM DTT, and the protein was eluted with the same buffer with additional 300 mM imidazole. The solution was further purified by injecting into Superdex 75 10/300 increase column (GE healthcare) equilibrated with buffer containing 20 mM Hepes pH 7.3, 500 mM NaCl, 2 mM DTT, and 5% glycerol. The purified ^DBDR^OCT4 was concentrated and stored at -80 ºC. The gene of the full-length OCT4 with an MBP-tag at N-terminus and a His-tag at C-terminus was inserted into the plasmid (pET30a). The plasmid was transformed into ArcticExpress (DE3) competent cells and grown in LB medium containing 20 μg/ml of gentamycin and 50 μg/ml of kanamycin at 30 ºC. When OD was about 0.8, 0.5 mM IPTG was added to induce protein expression at 10 ºC for at least 24 hrs. Protein purification followed the same procedure as ^MBP-DBDR^OCT4. The purified protein was concentrated to ∼24 µM and stored at -80 ºC.

### Preparation of DNA

DNA fragments were prepared by PCR amplification, followed by ethanol precipitation and purification using the POROS column. Briefly, the PCR products were pelleted by 75% ethanol containing 0.3 M NaAc at pH 5.2. The sample was incubated for 60 min at -20 ºC, followed by centrifugation. The pellet was resuspended in TE buffer. The sample was loaded to POROS column chromatography (GE Healthcare). The column was washed with buffer containing 20 mM Tris-HCl, pH 7.4, 5 mM 2-mercaptoethanol, and the DNA was eluted by a linear gradient of 0 to 2 M NaCl.

### Reconstitution of nucleosomes and nucleosome arrays

Purified recombinant histones in equal stoichiometric ratio were dissolved in ∼6 ml unfolding buffer (7 M guanidine-HCl, 20 mM Tris-Cl at pH 7.4, 10 mM DTT) and were dialyzed against refolding buffer (10 mM Tris-Cl at pH 7.4, 1 mM EDTA, 5 mM β-mercaptoethanol, 2 M NaCl, 0.1 mM PMSF) for about 12 hours. The mixture was centrifuged at 4000 rpm to remove any insoluble material. Soluble octamers were purified by size fractionation on a Superdex 200 gel filtration column. Purified histone octamers and DNA were mixed with a 1.3:1 ratio in high-salt buffer (2 M NaCl, 10 mM K/Na-phosphate at pH 7.4, 1 mM EDTA, 5 mM DTT). 1 ml mixture in a dialysis bag was placed in 600 mL of the high-salt buffer and dialyzed for 30 min followed by salt gradient dialysis. 3 L of a low-salt buffer (150 mM NaCl, 10 mM K/Na-phosphate at pH 7.4, 1 mM EDTA, 2 mM DTT) were gradually pumped into dialysis buffer with a flow rate of 2 ml/min for 18 h. The dialysis bag was then dialyzed against low-salt buffer for 30 min. The dialysis was done in the cold room. The sample was then incubated at 37 °C for 3-5 h. The mixture was centrifuged at 12,000 rpm to remove any insoluble material. The nucleosomes were further purified by ion-exchange chromatography (TSKgel DEAE, TOSOH Bioscience, Japan) to remove free DNA and histones. The purified nucleosomes were dialyzed against buffer containing 20 mM Hepes 7.3, 2 mM DTT.

To reconstitute 12 × 197 bp nucleosome array with a *LIN28B* nucleosome in the middle, or a ‘601’ nucleosome containing the Dde1 sites whose locations are the same as those in the *LIN28B* nucleosome, a pUC-18 plasmid was reconstructed with incorporation of sequentially ligated fragments of 5 × 197 bp W601 DNA, 1 × 197 bp *LIN28B* DNA or W601 DNA with Dde1 sites, and 6 × 197 bp W601 DNA. The resulting 12 × 197 bp DNA in the plasmid was verified by sequencing and restriction enzyme digestion. Large scale preparation and purification of the plasmid followed previous protocol (Zhou and Bai, 2021). Briefly, 12 × 197 bp DNA was released from the plasmid by EcoRV digestion (150 unit enzyme per 1 mg plasmid) and purified by stepwise PEG 6000 precipitation. The fractions containing the 12 × 197 bp DNA was used for nucleosome array reconstitution according to early methods (Dorigo et al., 2004). Saturation of nucleosome in the reconstituted 12 × 197 bp nucleosome array was verified by SmaI digestion, which showed predominant mono-nucleosome band on a 1.2% agarose gel. Linker histone H1.4-bound nucleosome array was made by mixing 1.3-fold of H1.4 (relative to molar concentration of nucleosomes in the array) with the nucleosome array in 10 mM Tris-HCl, pH 8.0, 1 mM EDTA, 1 mM DTT and 0.6 M NaCl buffer, followed by dialysis using the same buffer without NaCl.

### Cryo-EM sample preparation and Data collection

2 μM (50 μL) nucleosome containing human *LIN28B* DNA (162 bp), 6 μM (50 μL) scFv 60, and 40 μM (20 μL) ^MBP-DBDR^OCT4 were mixed in the buffer of 20 mM HEPES pH 7.3, 145 mM NaCl, 0.5 mM TCEP. The sample was incubated on ice for 0.5 h and then concentrated to ∼2 μM (nucleosome), which was loaded onto the glow-discharged holey carbon grid (Quantifoil 300 mesh Cu R1.2/1.3). All the grids were blotted for 3 s at 14 °C and 100% relative humidity using an FEI Vitrobot Mark IV plunger before being plunge-frozen in liquid nitrogen-cooled liquid ethane. For the native 162 bp *LIN28B* nucleosome sample without OCT4, and 187 bp site 0_mut free nucleosome, equal volume (25 μL) for each 4 μM nucleosome was mixed with 12 μM ScFv and the grids were prepared using the same parameter with nucleosome-^MBP-DBDR^OCT4 complex. Data were collected using a Titan Krios G3 electron microscope (Thermo-Fisher) operated at 300kV. Micrographs were acquired in super-resolution mode at the nominal magnification of 81,000x with 0.528 Å pixel size using a 20-eV slit post-GIF Gatan K3 camera. The dose rate on the camera was set to 15 e^-^/pixel/s. The total exposure time of each micrograph was 4 sec fractionated into 50 frames with 0.08 sec exposure time for each frame. Data collection were automated using the SerialEM software package (Mastronarde, 2005). A total of 7,800 micrographs were collected for the sample of the nucleosome bound to scFv and ^MBP-DBDR^OCT4, and a total of 2,446 micrographs were collected for the free *LIN28B* nucleosome sample. 8,235 micrographs were collected for 187 bp site 0_mut nucleosome sample.

### Image processing

All the datasets were processed using RELION/3.1.3 and CryoSPARC v3.2 following the standard procedures (Zivanov et al., 2018);(Punjani et al., 2017);(Zheng et al., 2017) (Figures S2 and S3). The averaged images without dose weighting were used for defocus determination using CTFFIND4.1 (Rohou and Grigorieff, 2015). Images with dose weighting were used for particle picking and extraction. Particles were automatically picked using Gautomatch (https://www.mrc-lmb.cam.ac.uk/kzhang/Gautomatch/). Bad particles were removed by 2D classification and 3D classification in RELION using 2x binned particles. One more round of 3D classification was made using finer angular sampling rate. Selected particles were submitted to Bayesian polishing and then imported to CryoSPARC. CTF-refinement and non-uniform refinement were made to generate the final maps for model building.

### Model building and structure analysis

For the free *LIN28B* nucleosome, an initial model of the nucleosome histone octamer and scFv was generated using the nucleosome structure (PDB: 7K61). They were fitted into the density map of the scFv-*LIN28B* nucleosome. The DNA was built into the map from scratch in COOT (Emsley et al., 2010). The histone octamer and scFv were optimized by manual rebuilding. The whole complex was refined using real space refinement in PHENIX (Adams et al., 2010). For the 187 bp site 0_mut nucleosome data set, the same procedure was used except the 162 bp *LIN28B* nucleosome structure was used as the initial model. For the complexes of the nucleosome bound to OCT4, the *LIN28B* nucleosome structure was used as an initial model of the nucleosome. Initial models of the POUS domain and DBDR of OCT4 were from the crystal structure of the mouse OCT4 DBDR bound to the DNA fragment (PDB: 3L1P). DNA was built into the map from scratch using COOT. Structure was optimized by manually rebuilding using COOT followed by further refinement using real space refinement in PHENIX. Figures were made using UCSF Chimera (Pettersen et al., 2004) and PyMOL (Version 1.8, Schrödinger, LLC. DeLano Scientific).

### Electrophoretic mobility shift assay

Nucleosomes at a final concentration of 0.3 μM were mixed with purified proteins. Typical binding reactions of complex formation were carried out for 30 min at room temperature in buffer containing 10 mM Hepes at pH 7.3, 60 mM NaCl, and 1 mM DTT. 10 μl of the binding reactions were analyzed on 5 % acrylamide gels in 0.2 x TBE at 120 V for 70 minutes at 4 °C. After electrophoresis, gels were stained with ethidium bromide (EtBr) and quantified using Image J. Binding data were fitted with the Hill equation and analyzed in Prism (Graphpad). For the competition reactions between H1.4 and ^MBP-DBDR^OCT4 for binding of the nucleosome, the H1.4 was mixed with the nucleosome at 1.6:1 ratio. The chaperone Nap1 at 2:1 ratio to the nucleosome was added to the mixture. The sample was incubated at room temperature for 15 min. ^DBDR^OCT4 was added and incubated at room temperature for 20 min. The final buffer contained 10 mM Hepes at pH 7.3, 2 mM DTT, 120 mM NaCl. 10 µl of the binding reaction solution was analyzed on 5 % acrylamide gels at 120 V for 70 min in 0.2 x TBE. Gels were stained with ethidium bromide (EtBr).

### Fluorescence resonance energy transfer assay

H1.4^K26C^ was labeled with Cy3 following the manufacture’s protocol (Cytiva). 1 mg H1.4 ^K26C^ was dissolved in 1 mL degassed buffer containing 20 mM HEPES 7.1, 500 mM NaCl and 1 mM TCEP. One vial of Cy3 maleimide was dissolved in dimethylformamide (DMF) and mixed with H1.4 ^K26C^. The mixture was agitated overnight at cold room. Size exclusive chromatography was used to separate the free Cy3 dye. H2A^L116C^ was labeled with Cy5 according to the previous publication (Shimko et al., 2011). 1 mg H2A^L116C^ was dissolved in 1 ml degassed buffer containing 800 mM HEPES 7.1, 1.5 mM Guanidine HCl, 1 mM TCEP. 1vial Cy5 maleimide (Cytiva, California) in DMF was mixed with H2A^L116C^ and incubated at room temperature for 5 hours. A Sephadex G-25 column was used to separate the free Cy5 dye. Fluorescent DNA fragments were produced using PCR by the Cy3 or Cy5 labeled primers (IDT). OCT4 was titrated into 200 nM-400 nM nucleosome in buffer 20 mM HEPES 7.1, 100mM NaCl, 0.5mM TCEP. In competition FRET assay, H1.4 was mixed with nucleosome with a ratio of 1.6:1 before addition of ^MBP-DBDR^OCT4. The chaperone Nap1 at 2:1 ratio to the nucleosome was added to prevent the non-specific binding. The fluorescence intensity was recorded using QuantaMaster (Photon Technology International). Excitation wavelength was set to 510 nm, and emission spectra were collected from 530nm to 730nm. Three independent experiments were performed.

### MNase and restriction enzyme digestion assay

50 µL of 1.2 μM *LIN28B* nucleosome was mixed with increasing amounts of ^MBP-DBDR^OCT4 in buffer (20 mM HEPES pH 7.3, 50 mM NaCl, 2 mM DTT). Then, 10 X MNase digestion buffer and 0.3 units of MNase (NEB) was added to each reaction. Samples were incubated at 37 °C for 30 min, and the enzyme was inactivated by adding 50 mM EGTA. Samples were then incubated with proteinase K for 1 hour at room temperature. DNA purification was made by mixing 25% (v/v) phenol–chloroform–isoamyl alcohol mixture (25:24:1, sigma). After centrifugation, the top solution was harvested. The final purified samples were loaded onto an 8 % acrylamide gels at 150 V for 70 minutes at 4 °C in 0.2 x TBE. Gels were stained with ethidium bromide (EtBr). For the ^DBDR^OCT4 facilitated digestion of nucleosome arrays assay, 150 ng of the nucleosome arrays were mixed with increasing amounts of ^DBDR^OCT4 in digestion buffer (10 ul, 10 mM Tris pH 8.0, 60 mM NaCl, 1 mM magnesium chloride, 2 mM DTT) at room temperature. 0.1 units restriction enzyme DdeI (NEB), or 0.1 units restriction enzyme BanI (NEB), or 0.5 units restriction enzyme Sau96I (NEB) was added to each reaction solution. Samples were incubated at 37 °C for 40 min, and the enzyme was inactivated by incubating at 65 °C for 20 min. Samples were then incubated with proteinase K at 50 °C for 60 min. DNA was purified by mixing the sample with 25 % (v/v) phenol–chloroform–isoamyl alcohol mixture (25:24:1, sigma). After centrifugation, the top solution was harvested. The final purified samples were loaded onto a 1 % agarose gel stained with SYBR Safe dye (Invitrogen). Electrophoresis was performed at 150 V in 0.2 x TBE buffer for 30 min. Three independent experiments were performed. Band intensities for the digestion product and input were measured using ImageJ to calculate digestion efficiency.

## Acknowledgments

We thank Dr. Bingquan Gao for constructing the ^MBP-DBDR^OCT4 plasmid, Dr. Rick Huang and Ms. Allison Zeher for assistance in cryo-EM Data collection, Dr. Kenneth Zaret and Mr. Jacob Licht for helpful comments on the manuscript. The cryo-EM work utilized NCI-NIH IRP Cryo-EM Consortium (NICE) microscopy resource and NIH high performance computing Biowulf system for data processing. This work is supported by the intramural research program at the Center for Cancer Research, National Cancer Institute, National Institutes of Health.

## Author contributions

Y.B. conceived the project. R.G. and T.L. conducted the experiment. B-R. Z. provided the scFv and H1.4 proteins and reconstituted the nucleosome array. Y.B., R.G., and T.L. analyzed the structure and wrote the paper.

