## Supplemental Figures and Tables for "Structural mechanism of *LIN28B* nucleosome targeting by OCT4 for pluripotency"

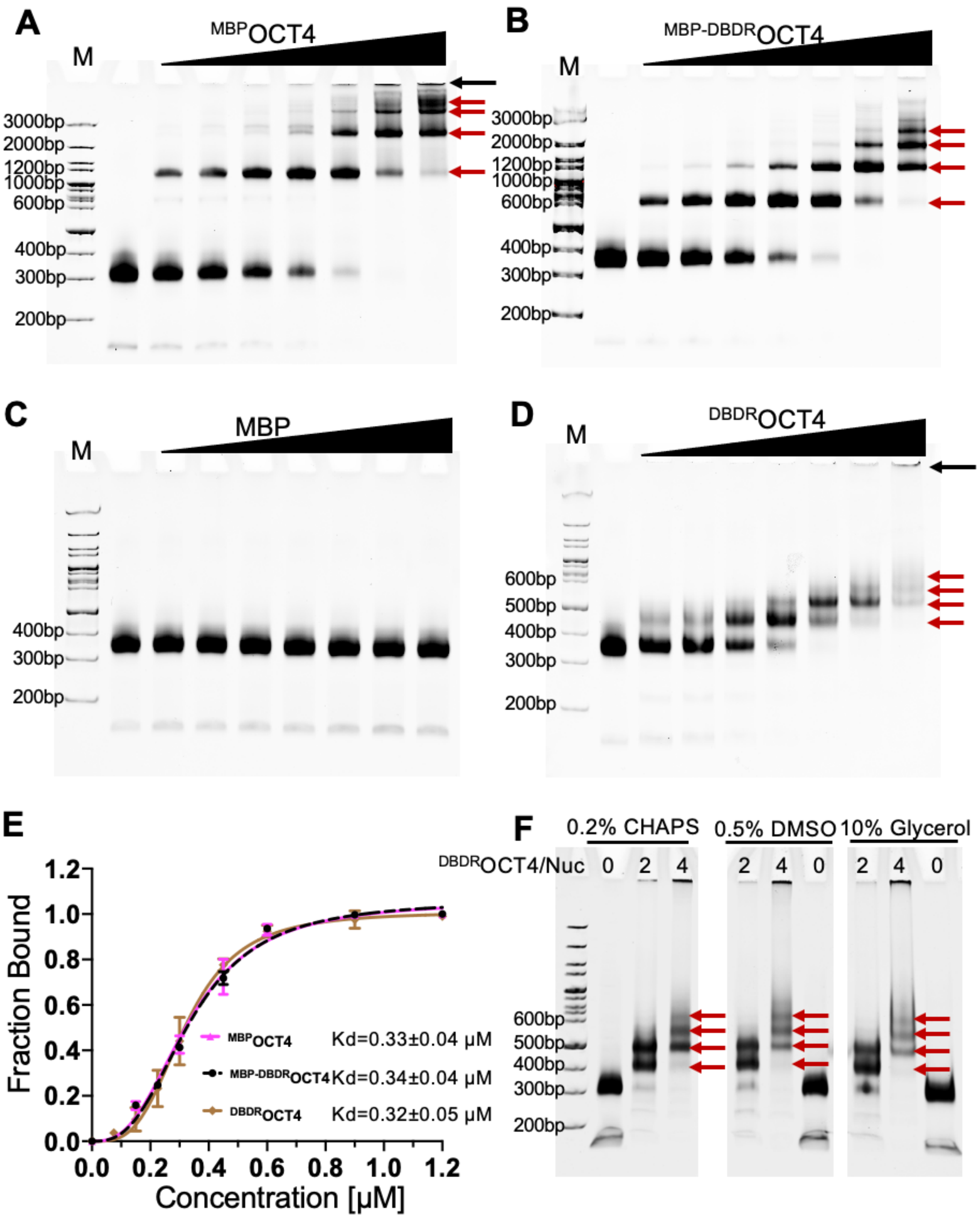


**Figure S1. Characterization of OCT4 binding to human *LIN28B* nucleosome, related to Figure 1 and Figure 2**

(A) EMSA of nucleosome association with $^{MBP}$OCT4. The nucleosome concentration is 0.3 μM. The ratios of the proteins over the nucleosome are 0, 0.5, 0.75, 1.0, 1.5, 2.0, 3.0, and 4.0, respectively. The same concentrations are used in (B-D below).

(B) EMSA of nucleosome association with $^{MBP\text{-}DBDR}$OCT4.

(C) EMSA of nucleosome association with MBP.

(D) EMSA of nucleosome association with $^{\mathrm{DBDR}}$OCT4.

(E) Quantification of the *LIN28B* nucleosome binding by $^{\mathrm{MBP}}$OCT4, $^{\mathrm{MBP\text{-}DBDR}}$OCT4, and $^{\mathrm{DBDR}}$OCT4. The apparent Kd values were obtained by fitting the data to the Hill equation. The Hill coefficient is ~3. Error bars represent standard deviation values from at least three independent experiments.

(F) EMSA showing the buffers that improve $^{\mathrm{DBDR}}$OCT4-nucleosome solubility. “M” stands for DNA marker. The black arrow indicates aggregation. The red arrows highlight the formation of multiple complexes.

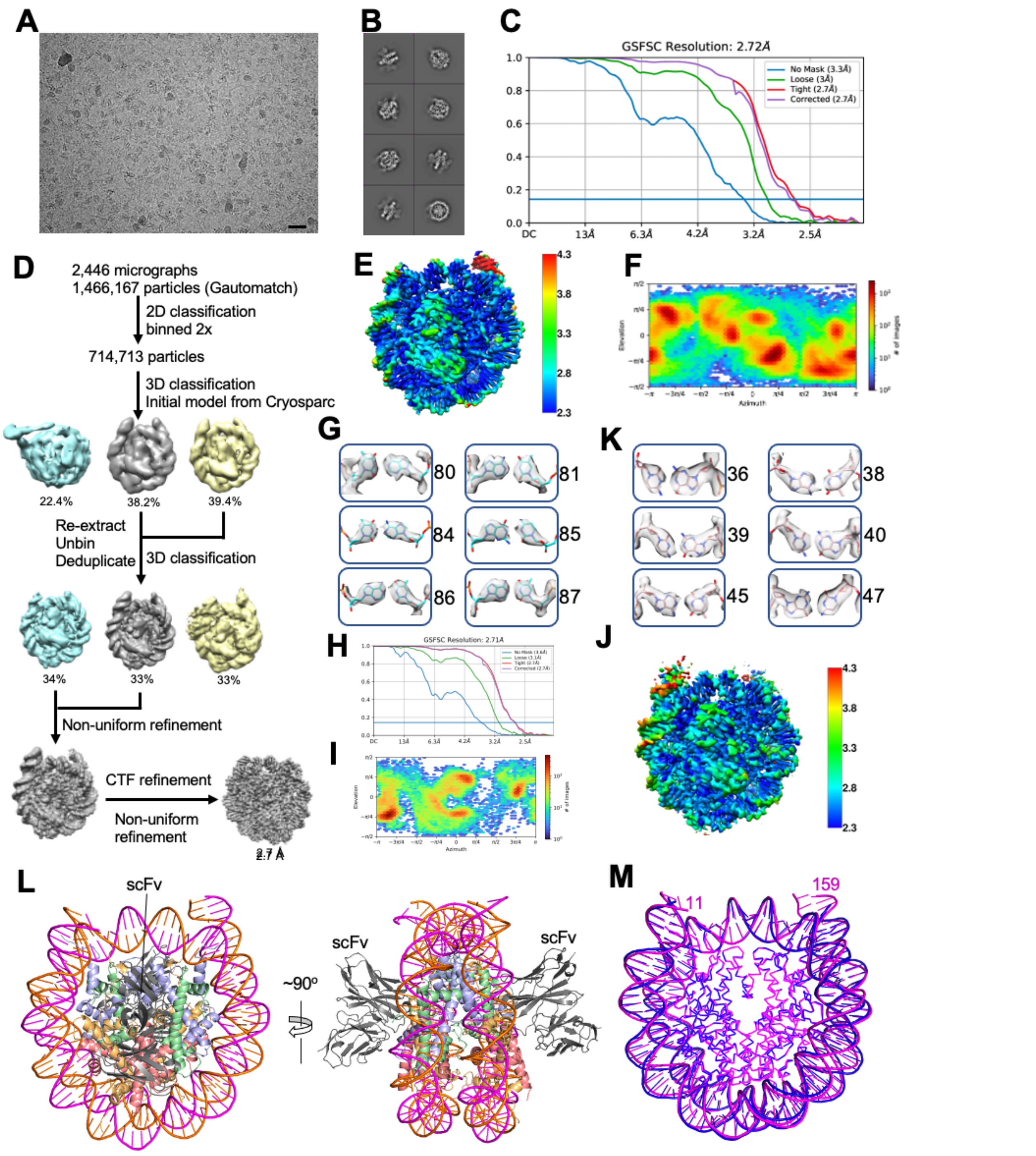


**Figure S2. Data collection and processing for human *LIN28B* nucleosomes, related to Figure 1 and Figure 5**

(A) Raw image of the 162 *LIN28B* nucleosome and scFv complex. (B-G) below are related to this data set.
(B) 2D classification results.
(C) FSC curves.

(D) Particle analysis.
(E) Local resolution.
(F) Particle orientation distribution.
(G) Illustration of density maps for the representative base pairs.
(H) FSC curves for the 187 bp *LIN28B* nucleosome with site 0 mutation bound to scFv.
(I) Particle orientation distribution for the 187 bp *LIN28B* nucleosome with site 0 mutation bound to scFv.
(J) Local resolution for the 187 bp *LIN28B* nucleosome with site 0 mutation bound to scFv.
(K) Illustration of density maps for the representative base pairs for the 187 bp *LIN28B* nucleosome with site 0 mutation bound to scFv.
(L) Structure of the 162 bp *LIN28B* scFv-nucleosome complex, highlighting the location of scFv (grey).
(M) Structure of 162 bp *LIN28B* nucleosome (magenta) aligned with the crystal structure of the 145 bp '601' nucleosome core particle (PDB ID: 2NZD; blue). DNAs are shown with cartoon. Histones are shown with C$\alpha$ traces.

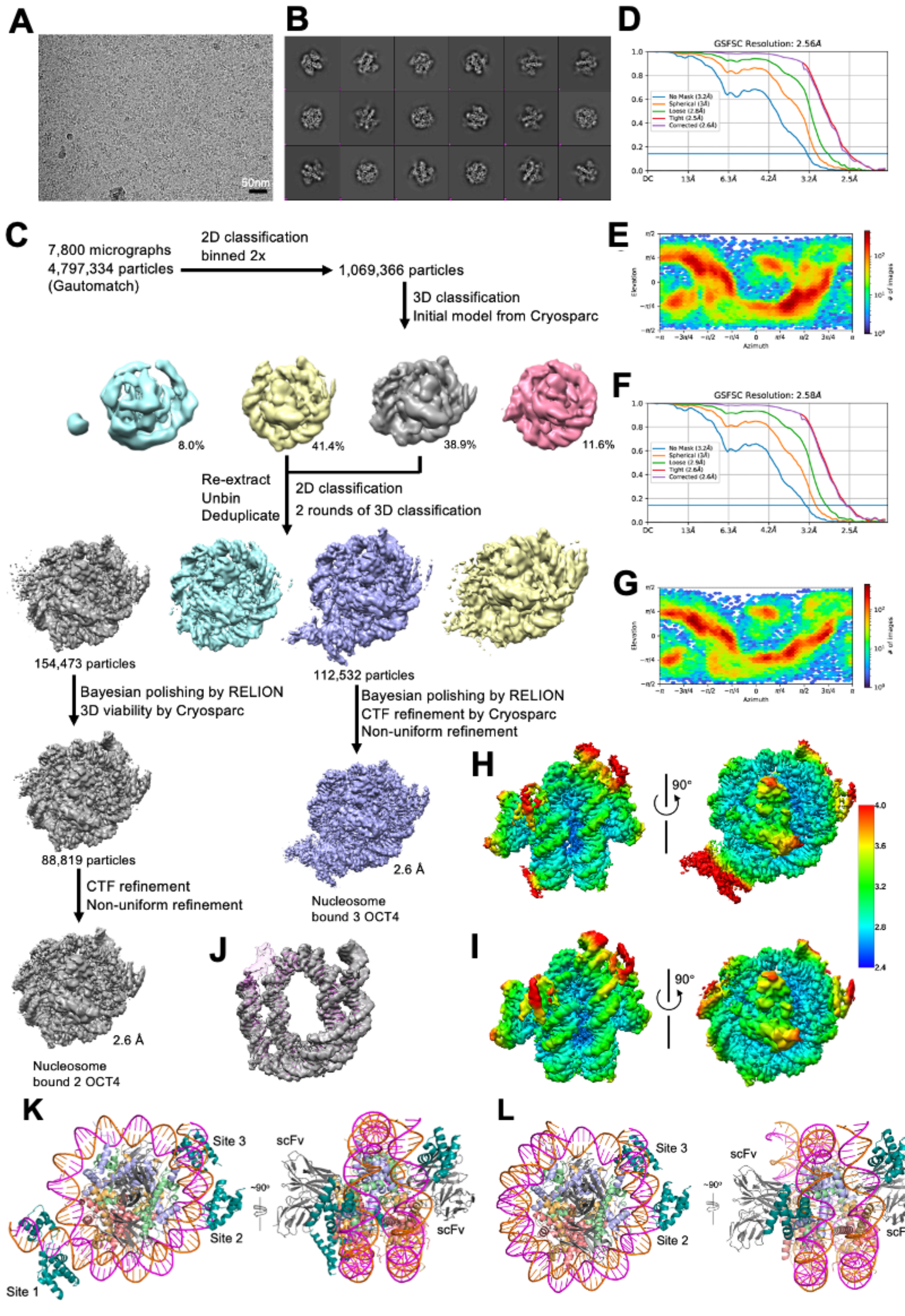


**Figure S3. Data collection and processing for human *LIN28B* nucleosome bound to $^{MBP\text{-}DBDR}$OCT4, related to Figure 1 and Figure 4**

(A) Raw image.

(B) 2D classification results.
(C) Particle analysis.
(D) FSC curve for the nucleosome bound to three $^{\text{MBP-DBDR}}$OCT4s.
(E) Particle orientation distribution for the nucleosome bound to three $^{\text{MBP-DBDR}}$OCT4s.
(F) FSC curve for the nucleosome bound to two $^{\text{MBP-DBDR}}$OCT4s.
(G) Particle orientation distribution for the nucleosome bound to two $^{\text{MBP-DBDR}}$OCT4s.
(H) Local resolution for the nucleosomes bound to three $^{\text{MBP-DBDR}}$OCT4.
(I) Local resolution for the nucleosomes bound to two $^{\text{MBP-DBDR}}$OCT4s. MBP was not observed, likely caused by the flexible linker and small size.
(J) Overlay of the density maps of the free 162 bp *LIN28B* nucleosome (magenta) and the nucleosome bound to two OCT4s (gray).
(K) Three $^{\text{MBP-DBDR}}$OCT4 molecules bind the 162 bp *LIN28B* nucleosome.
(L) Two $^{\text{MBP-DBDR}}$OCT4 molecules bind the 162 bp *LIN28B* nucleosome. In these structures, scFv binds to the core histones, forming no direct contacts with DNA and $^{\text{MBP-DBDR}}$OCT4.

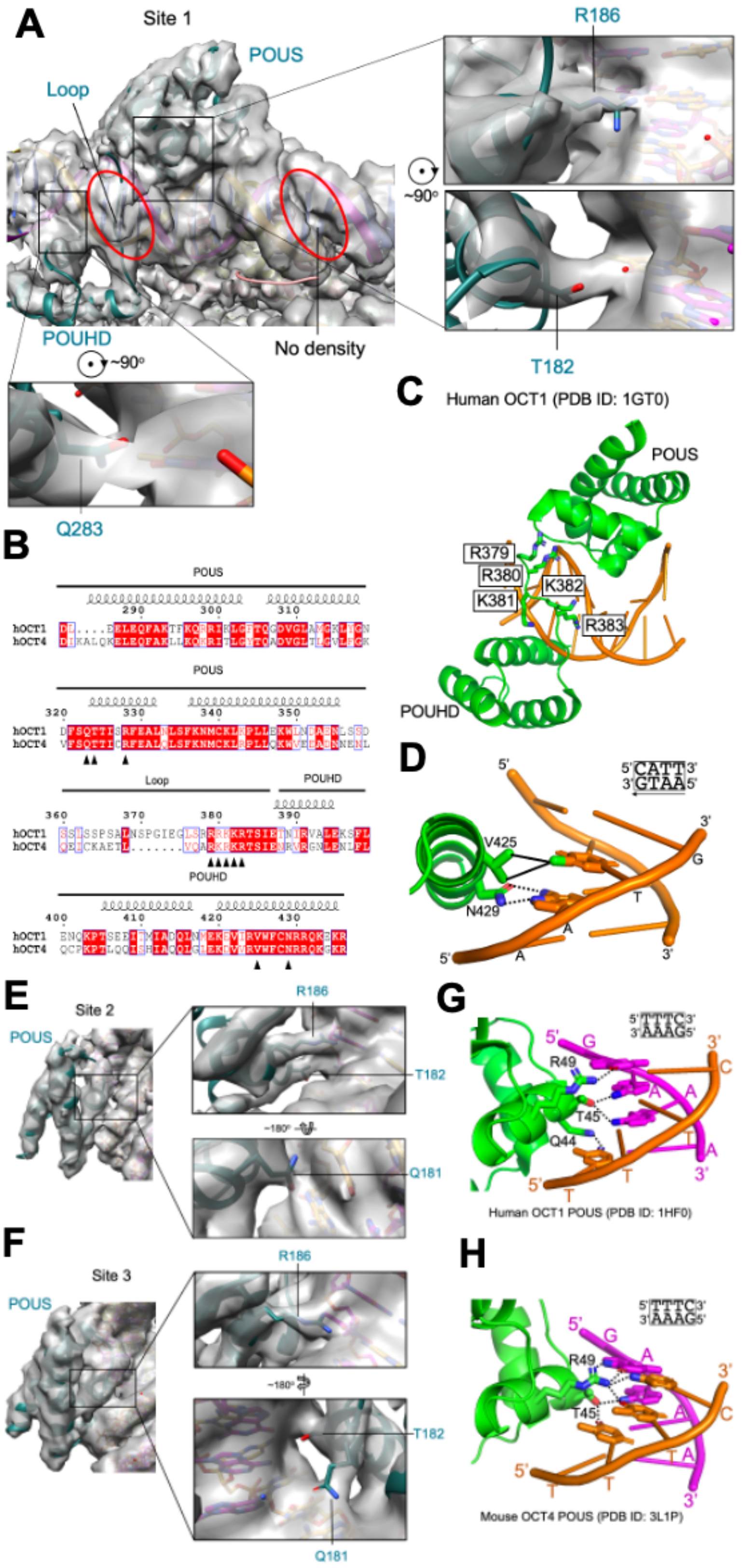


**Figure S4. The density maps at the interface between $^{\text{MBP-DBDR}}$OCT4 and the nucleosome and formation of hydrogen bonds, related to Figure 3**

(A) Identification of densities for the side chains of the residues that form hydrogen bonds with DNA at site 1. The open red ovals highlight the difference in densities at the two neighbor minor grooves. The extra density at site 1 reveals the interactions between basic patch residues and the DNA minor groove.
(B) Sequence alignment of DBDRs of human OCT4 and OCT1.
(C) Basic patch residues in the loop between POUS and POUHD interact with the minor groove of DNA (orange).
(D) Hydrogen bonds and hydrophobic interactions between POUHD and the noncanonical DNA motif (AATG) in the OCT1-DNA complex. OCT1 POUHD in green and DNA in orange.
(E) Density map at site 2.
(F) Density map at site 3.
(G) Human OCT1 POUS bound to noncanonical motif TTTC (Reményi et al., 2001).
(H) Mouse OCT4 POUS bound to noncanonical motif TTTC (Esch et al., 2013).

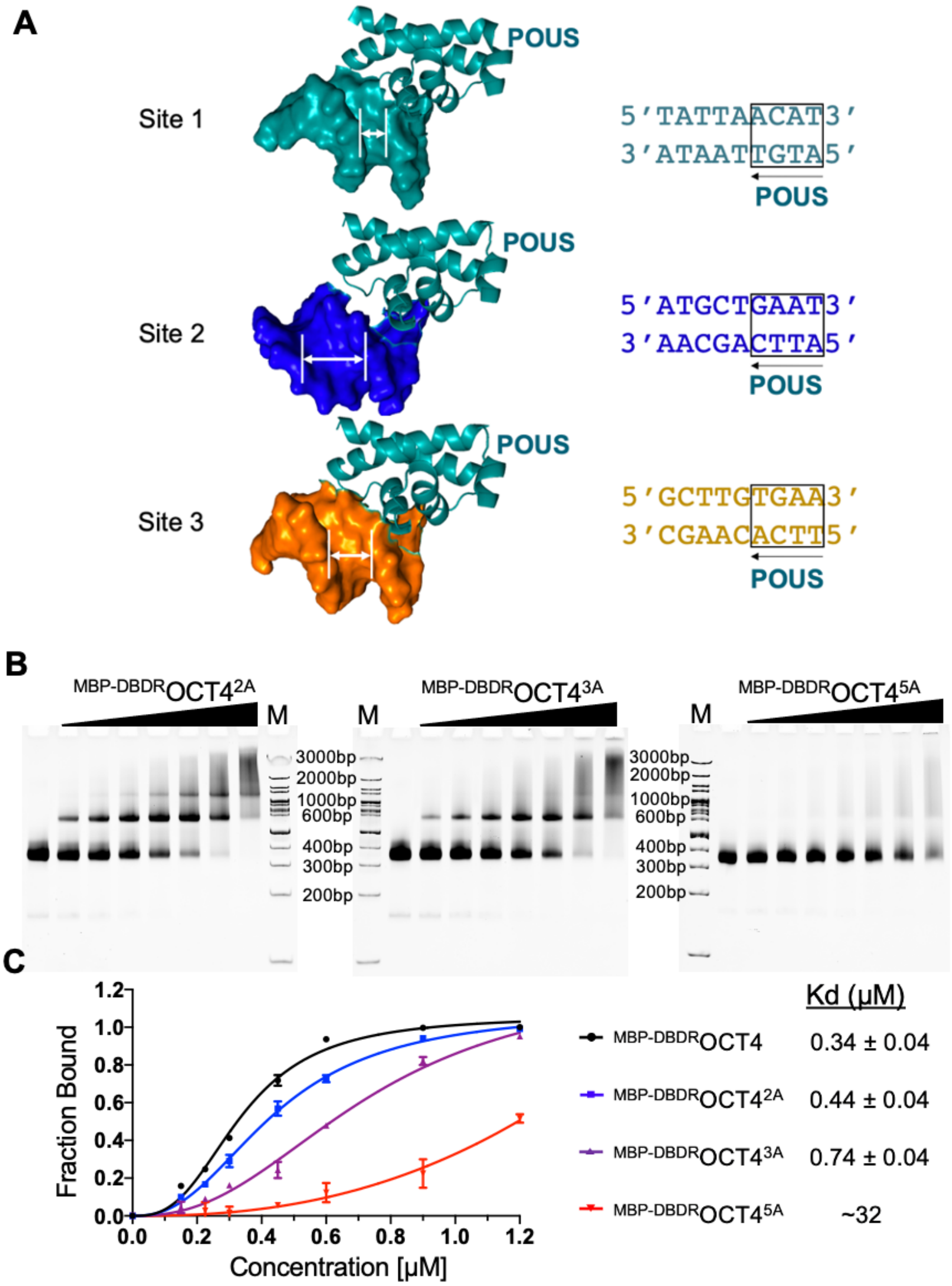


**Figure S5. The role of $^{DBDR}$OCT4 loop basic patch in Nu-motif binding, related to Figure 3**

(A) Illustration of the narrow minor groove width at site 1 with an AT-track near the POUS domain and lack of AT-tracks at sites 2 and 3.

(B) EMSA experiments for OCT4 mutation R232A/R234A (left), R232A/K233A/R234A (middle), and R230A/K231A/R232A/K233A/R234A, repectively.

(C) Quantification of the of gel bands and fitting of the data to the Hill equation to obtain apparent Kd values (right). The WT data were from Extended Data Figure.1. For these experiments, the nucleosome concentration is 0.3 μM. The ratios of the proteins over the nucleosome are 0, 0.5, 0.75, 1.0, 1.5, 2.0, 3.0, and 4.0, respectively. Error bars represent standard deviation values from three independent experiments.

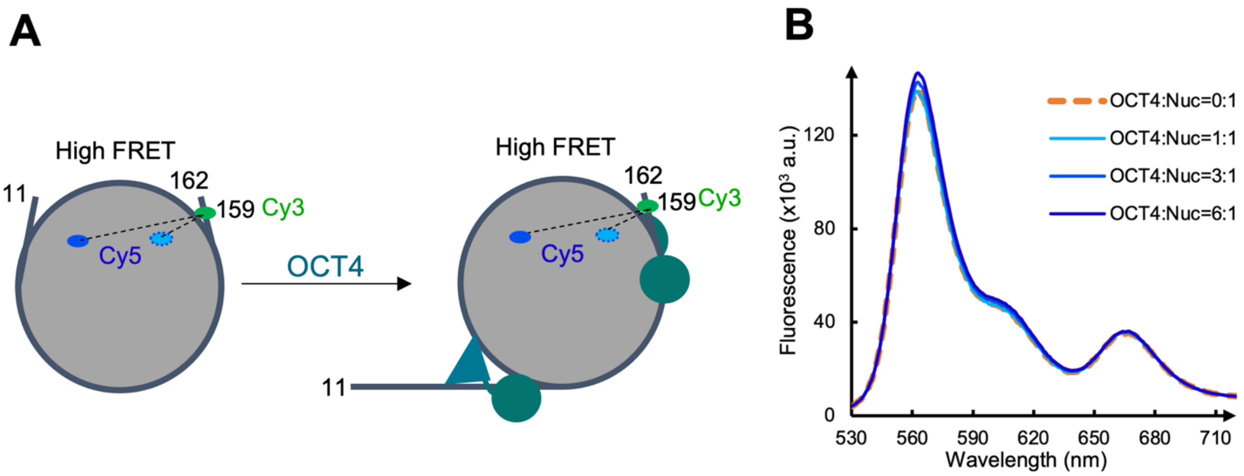


**Figure S6. FRET assay of OCT4 binding effect on exit DNA, related to Figure 5**

(A) Diagram illustration of Cy5 and Cy3 labeling on H2A and the DNA at the exit site, respectively (Left) and expected FRET signals (no change) with binding of OCT4 to the nucleosome.

(B) Measured FRET signals with increase of $^{MBP\text{-}DBDR}$OCT4 concentration.

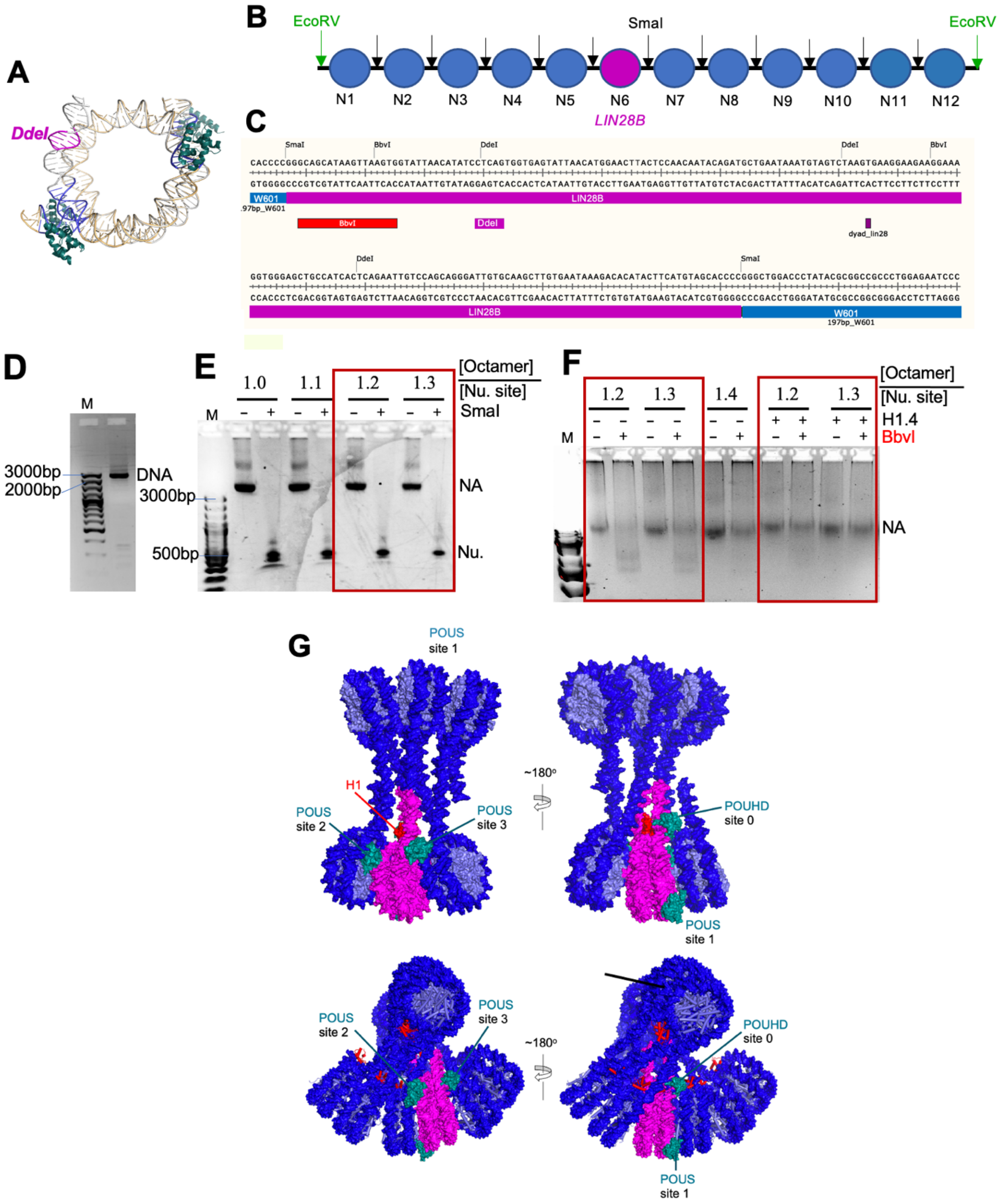


**Figure S7. OCT4 opens the H1-condensed nucleosome array, related to Figure 6 and Figure 7**

(A) Illustration of the DNA cutting site in the *LIN28B* nucleosome.

(B) Diagram illustration of the design of nucleosome array consisting of the *LIN28B* nucleosome.

(C) The *LIN28B* DNA sequence in the nucleosome array.
(D) Agarose gel showing purified 12 x 197 bp DNA.
(E) Agarose gel showing reconstituted nucleosome arrays (NA) with different ratios of core histone octamer over nucleosome sites. The red box highlights the saturated nucleosome arrays that led to mononucleosomes (Nu.) when cut by SmaI enzyme.
(F) Agarose gel showing reconstituted nucleosome arrays with different ratios of core histone octamer over the nucleosome site in the DNA with and without H1.4 and BbvI enzyme. BbvI is specific to the expanded *LIN28B* DNA. The red boxes highlight the reconstituted H1.4-condnesed nucleosome array that is resistant to the digestion by BbvI relative to the nucleosome array without H1.4.
(G) Structural modeling showing clashes of POUS and POUHD bound to their Nu-motifs in the *LIN28B* chromatosome with the neighboring chromatosomes in the ladder-like (top) (Garcia-Saez et al., 2018) and twisted double-helix (bottom) (Song et al., 2014) nucleosome array structures.

### Table S1. Data collection and model refinement parameters.

| **Data collection and processing** | |
|---|---|
| Magnification | 81,000 |
| Voltage (kV) | 300 |
| Exposure time (s/frame) | 0.08 |
| Number of frames | 50 |
| Electron exposure (e-/Å$^2$) | 53.8 |
| Defocus range (μm) | -1.0 ~ -2.0 |
| Pixel size (Å) | 0.528 |
| Symmetry imposed | C1 |

| | ***LIN28B* Nuc-ScFv$_2$ PDB ID: 7U0J EMDB ID: 26261** | **187bp site0_mut *LIN28B* Nuc-ScFv$_2$ PDB ID: 8DK5 EMDB ID: 27483** | **Nuc-scFv$_2$-($^{MBP\text{-}DBDR}$OCT4)$_2$ PDB ID: 7U0I EMDB ID: 26260** | **Nuc-scFv$_2$-($^{MBP\text{-}DBDR}$OCT4)$_3$ PDB ID: 7U0G EMDB ID: 26258** |
|---|---|---|---|---|
| Initial particle images (no.) | 1,466,167 | 1,813,056 | 1,069,366 | 1,069,366 |
| Final particle images (no.) | 359,203 | 49,406 | 88,819 | 112,532 |
| Map resolution (Å) | 2.7 | 2.7 | 2.6 | 2.6 |
| FSC threshold | 0.143 | 0.143 | 0.143 | 0.143 |
| Map resolution range | 2.3-8.4 | 2.3-11.1 | 2.4-7.1 | 2.4-9.1 |
| Map sharpening $B$ factor (Å$^2$) | -86 | -80 | -20 | -20 |
| **Refinement** | | | | |
| Non-hydrogen atoms | 15,622 | 15,621 | 16,888 | 17,327 |
| Protein residues | 1,210 | 1,210 | 1,368 | 1,510 |
| Nucleotide | 298 | 298 | 298 | 266 |
| ***B* factors (Å$^2$)** | | | | |
| Protein | 62.25 | 61.98 | 79.43 | 97.67 |
| Nucleic acids | 98.79 | 82.06 | 120.09 | 110.25 |
| **R.m.s deviations** | | | | |
| Bond lengths (Å) | 0.008 | 0.011 | 0.010 | 0.006 |
| Bond angles (°) | 0.930 | 1.114 | 0.862 | 0.777 |
| **Validation** | | | | |
| MolProbity score | 1.66 | 1.83 | 2.09 | 1.94 |
| Clashscore | 6.00 | 9.46 | 14.30 | 14.52 |
| Poor rotamers (%) | 0.1 | 0.1 | 0.2 | 0.1 |
| Ramachandran Plot | | | | |
| Favored (%) | 95.28 | 95.24 | 95.98 | 96.14 |
| Allowed (%) | 4.64 | 4.64 | 3.88 | 3.79 |
| Disallowed (%) | 0.08 | 0.08 | 0.14 | 0.07 |

### Table S2. A list of protein and DNA sequences and primers

| | |
|---|---|
| MBP-DBDR OCT4 Protein with proteinase cutting site underlined | MKEAKIEEGKLVIWINGDKGYNGLAEVGKKFEKDTGIKVTVEHPDK<br>LEEKFPQVAATGDGPDIIFWAHDRFGGYAQSGLLAEITPDKAFQDKL<br>YPFTWDAVRYNGKLIAYPIAVEALSLIYNKDLLPNPPKTWEEIPALD<br>KELKAKGKSALMFNLQEPYFTWPLIAADGGYAFKYENGKYDIKDV<br>GVDNAGAKAGLTFLVDLIKNKHMNADTDYSIAEAAFNKGETAMTI<br>NGPWAWSNIDTSKVNYGVTVLPXFKGQPSKPFVGVLSAGINAASPN<br>KELAKEFLENYLLTDEGLEAVNKDKPLGAVALKSYEEELAKDPRIA<br>ATMENAQKGEIMPNIPQMSAFWYAVRTAVINAASGRQTVDAALAA<br>AQTNAGSENLYFQGSVDSAAASDIKALQKELEQFAKLLKQKRITLG<br>YTQADVGLTLGVLFGKVFSQTTICRFEALQLSFKNMCKLRPLLQKW<br>VEEADNNENLQEICKAETLVQARKRKRTSIENRVRGNLENLFLQCPK<br>PTLQQISHIAQQLGLEKDVVRVWFCNRRQKGKRSSSEFHHHHHH |
| R232A/R234A_F | CAGAAACCCTCGTGCAGGCCCGAAAGGCAAAGGCA<br>ACCAGTATCGAGAACCGAG |
| R232A/R234A_R | CTCGGTTCTCGATACTGGTTGCCTTTGCCTTTCGGGCCTGCACGAG<br>GGTTTCTG |
| R232A/K233A/R234A_F | CAGAAACCCTCGTGCAGGCCCGAAAGGCAGCGGCA<br>ACCAGTATCGAGAACCGAG |
| R232A/K233A/R234A-R | CTCGGTTCTCGATACTGGTTGCCGCTGCCTTTCGGGCCTGCACGA<br>GGGTTTCTG |
| R230A/K231A/R232A/K233A/R234A _F | TGCAGGCGGCGGCTGCAGCGGCCACCAGTATCGAGAACCGAGT |
| R230A/K231A/R232A/K233A/R234A _R | ATACTGGTGGCCGCTGCAGCCGCCGCCTGCACCAGGGTTTC |
| 162_*LIN28B*_DNA | AGTGGTATTAACATATCCTCAGTGGTGAGTATTAACATGGAACTT<br>ACTCCAACAATACAGATGCTGAATAAATGTAGTCTAAGTGAAGG<br>AAGAAGGAAAGGTGGGAGCTGCCATCACTCAGAATTGTCCAGCA<br>GGGATTGTGCAAGCTTGTGAATAAAGACA |
| 162_*LIN28B*_DNA_template_F | AGTGGTATTAACATATCCTCAGTGGTGAGTATTAACATGGAACTT<br>ACTCCAACAATACAGATGCTGAATAAATGTAGTCTAAGTGAAGG<br>AAGAAGG |
| 162_*LIN28B*_DNA_template_R | TGTCTTTATTCACAAGCTTGCACAATCCCTGCTGGACAATTCTGA<br>GTGATGGCAGCTCCCACCTTTCCTTCTTCCTTCACTTAGACTACAT<br>TTATT |
| 162_*LIN28B*_DNA_F | AGTGGTATTAACATATCCTC |
| 162_*LIN28B*_DNA_R | TGTCTTTATTCACAAGCTTG |
| 187_*LIN28B*_DNA | GCATAAGTTAAGTGGTATTAACATATCCTCAGTGGTGAGTATTAA<br>CATGGAACTTACTCCAACAATACAGATGCTGAATAAATGTAGTCT<br>AAGTGAAGGAAGAAGGAAAGGTGGGAGCTGCCATCACTCAGAAT<br>TGTCCAGCAGGGATTGTGCAAGCTTGTGAATAAAGACACATACTT<br>CATGTAGT |
| 187_*LIN28B*_DNA_F | GCATAAGTTAAGTGGTATTAACATATCCTCAGTGG |
| 187_*LIN28B*_DNA_R | ACTACATGAAGTATGTGTCTTTATTCACAAGC |
| 187_*LIN28B*_site 0 mut_DNA_template_F | AGTGGAGAGAAAGAATCCTCAGTGGTGAGTATTAACATGGAACT<br>TACTCCAACAATACAGATGCTGAATAAATGTAGTCTAAGTGAAG<br>GAAGAAGG |
| 187_*LIN28B*_site 0 mut_DNA_F | GCATAAGTTAAGTGGAGAGAAAGAATCCTCAGTGG |
| Cy3_*LIN28B*_11F | /5Cy3/ACATATCCTCAGTGGTGAGTATTAACATGG |
| Cy3_*LIN28B*_159R | /5Cy3/CTTTATTCACAAGCTTGCACAATCC |
| 601_3sites_ template F | ATCGAGAATCCCGGTGCAGTATTAACATGGTTGGTCGTAGACAGC<br>TCTAGCAC CTGAATAACGCACGTACGCGCTGTCCCCCGCGTTTTA |
| 601_3sites_ template R | ATATTCACAATATATCTGACACGTGCCTGGAGACTAGGGAGTAAT<br>CCCCTTGGCGGTTAAAACGCGGGGGACAGCGCGTACGTGCGTTAT<br>TCAG |

| | |
|---|---|
| 601_3sites_F | ATCGAGAATCCCGGTGCAGTA |
| 601_3sites_R | ATATTCACAATATATCTGACACGTGCCTGGAG |
| 12x197 bp DNA (197 bp extended native *LIN28B* DNA underlined) sequence | ATCCCTCCACCAGGGGGGCCCCCCGGGCTGGACCCTATACGCGGC<br>CGCCCTGGAGAATCCCGGTGCCGAGGCCGCTCAATTGGTCGTAGA<br>CAGCTCTAGCACCGCTTAAACGCACGTACGCGCTGTCCCCCGCGT<br>TTTAACCGCCAAGGGGATTACTCCCTAGTCTCCAGGCACGTGTCA<br>GATATATACATCCTGTGCATGTATTGAACAGCGACCACCCCGGGC<br>TGGACCCTATACGCGGCCGCCCTGGAGAATCCCGGTGCCGAGGC<br>CGCTCAATTGGTCGTAGACAGCTCTAGCACCGCTTAAACGCACGT<br>ACGCGCTGTCCCCCGCGTTTTAACCGCCAAGGGGATTACTCCCTA<br>GTCTCCAGGCACGTGTCAGATATATACATCCTGTGCATGTATTGA<br>ACAGCGACCACCCCGGGCTGGACCCTATACGCGGCCGCCCTGGA<br>GAATCCCGGTGCCGAGGCCGCTCAATTGGTCGTAGACAGCTCTAG<br>CACCGCTTAAACGCACGTACGCGCTGTCCCCCGCGTTTTAACCGC<br>CAAGGGGATTACTCCCTAGTCTCCAGGCACGTGTCAGATATATAC<br>ATCCTGTGCATGTATTGAACAGCGACCACCCCGGGCTGGACCCTA<br>TACGCGGCCGCCCTGGAGAATCCCGGTGCCGAGGCCGCTCAATTG<br>GTCGTAGACAGCTCTAGCACCGCTTAAACGCACGTACGCGCTGTC<br>CCCCGCGTTTTAACCGCCAAGGGGATTACTCCCTAGTCTCCAGGC<br>ACGTGTCAGATATATACATCCTGTGCATGTATTGAACAGCGACCA<br>CCCCGGGCTGGACCCTATACGCGGCCGCCCTGGAGAATCCCGGTG<br>CCGAGGCCGCTCAATTGGTCGTAGACAGCTCTAGCACCGCTTAAA<br>CGCACGTACGCGCTGTCCCCCGCGTTTTAACCGCCAAGGGGATTA<br>CTCCCTAGTCTCCAGGCACGTGTCAGATATATACATCCTGTGCAT<br>GTATTGAACAGCGACCACCCCGGGCAGCATAAGTTAAGTGGTATT<br>AACATATCCTCAGTGGTGAGTATTAACATGGAACTTACTCCAACA<br>ATACAGATGCTGAATAAATGTAGTCTAAGTGAAGGAAGAAGGAA<br>AGGTGGGAGCTGCCATCACTCAGAATTGTCCAGCAGGGATTGTGC<br>AAGCTTGTGAATAAAGACACATACTTCATGTAGCACCCCGGGCTG<br>GACCCTATACGCGGCCGCCCTGGAGAATCCCGGTGCCGAGGCCG<br>CTCAATTGGTCGTAGACAGCTCTAGCACCGCTTAAACGCACGTAC<br>GCGCTGTCCCCCGCGTTTTAACCGCCAAGGGGATTACTCCCTAGT<br>CTCCAGGCACGTGTCAGATATATACATCCTGTGCATGTATTGAAC<br>AGCGACCACCCCGGGCTGGACCCTATACGCGGCCGCCCTGGAGA<br>ATCCCGGTGCCGAGGCCGCTCAATTGGTCGTAGACAGCTCTAGCA<br>CCGCTTAAACGCACGTACGCGCTGTCCCCCGCGTTTTAACCGCCA<br>AGGGGATTACTCCCTAGTCTCCAGGCACGTGTCAGATATATACAT<br>CCTGTGCATGTATTGAACAGCGACCACCCCGGGCTGGACCCTATA<br>CGCGGCCGCCCTGGAGAATCCCGGTGCCGAGGCCGCTCAATTGGT<br>CGTAGACAGCTCTAGCACCGCTTAAACGCACGTACGCGCTGTCCC<br>CCGCGTTTTAACCGCCAAGGGGATTACTCCCTAGTCTCCAGGCAC<br>GTGTCAGATATATACATCCTGTGCATGTATTGAACAGCGACCACC<br>CCGGGCTGGACCCTATACGCGGCCGCCCTGGAGAATCCCGGTGCC<br>GAGGCCGCTCAATTGGTCGTAGACAGCTCTAGCACCGCTTAAACG<br>CACGTACGCGCTGTCCCCCGCGTTTTAACCGCCAAGGGGATTACT<br>CCCTAGTCTCCAGGCACGTGTCAGATATATACATCCTGTGCATGT<br>ATTGAACAGCGACCACCCCGGGCTGGACCCTATACGCGGCCGCC<br>CTGGAGAATCCCGGTGCCGAGGCCGCTCAATTGGTCGTAGACAG<br>CTCTAGCACCGCTTAAACGCACGTACGCGCTGTCCCCCGCGTTTT<br>AACCGCCAAGGGGATTACTCCCTAGTCTCCAGGCACGTGTCAGAT<br>ATATACATCCTGTGCATGTATTGAACAGCGACCACCCCGGGCTGG<br>ACCCTATACGCGGCCGCCCTGGAGAATCCCGGTGCCGAGGCCGCT<br>CAATTGGTCGTAGACAGCTCTAGCACCGCTTAAACGCACGTACGC<br>GCTGTCCCCCGCGTTTTAACCGCCAAGGGGATTACTCCCTAGTCT<br>CCAGGCACGTGTCAGATATATACATCCTGTGCATGTATTGAACAG<br>CGACCACCCCGTGGGCCACAAAAGTGGGAT |
